# A high confidence *Physcomitrium patens* plasmodesmata proteome by iterative scoring and validation reveals diversification of cell wall proteins during evolution

**DOI:** 10.1101/2022.06.01.492581

**Authors:** Sven Gombos, Manuel Miras, Vicky Howe, Lin Xi, Mathieu Pottier, Neda S. Kazemein Jasemi, Moritz Schladt, Jona Ejike, Ulla Neumann, Sebastian Hänsch, Franziska Kuttig, Zhaoxia Zhang, Marcel Dickmanns, Peng Xu, Torsten Stefan, Wolfgang Baumeister, Wolf B. Frommer, Rüdiger Simon, Waltraud X. Schulze

## Abstract

Cells of multicellular organisms exchange nutrients, building blocks and information. In animals, this happens via gap junctions, in plants via plasmodesmata (PD). PD have striking properties, translocating a large range of molecules from ions, to metabolites, RNA and proteins up to 40 kDa. PD are hard to characterize due to being deeply embedded into cell walls and the presence of several membranes. While previous studies of protein composition of PD from angiosperms identified large lists of proteins, few were validated. Here, we developed a PD scoring approach in conjunction with systematic localization on a large scale to define a high-confidence PD proteome of *Physcomitrium patens*. This high confidence PD proteome comprises nearly 300 proteins, which together with the *bona fide* PD proteins from literature, are made available in the public PDDB database. Conservation of localization across plant species strengthens the reliability of plant PD proteomes and provides a basis for exploring the evolution of this important organelle. In particular, the *P. patens* PD proteome was highly enriched in cell wall modifying proteins. Callose-degrading glycolyl hydrolase family 17 (GHL17) proteins are presented as an abundant PD protein family with representatives across an evolutionary scale. Exclusively members of the alpha-clade of the GHL17 family are shown to be PD localized and their orthologs occur only in plant species which have developed PD. Members of the EXORDIUM-family and xyloglucan transglycosylases are additional cell-wall located proteins highly abundant in the *P. patens* PD proteome also showing evolutionary diversification of PD localized family members from other clade members.

## Introduction

Plasmodesmata (PD) are important and complex structures present in all land plants. During evolution of multicellular life forms, multicellular complexity from single cells during development had to be achieved, and exchange of nutrients and information between was required to coordinate development and distinct cellular differentiation. The cellular structures and mechanisms facilitating intracellular communication evolved along different pathways. In animals, different types of gap junctions mediate intercellular exchange of small molecules, while primary cilia evolved to coordinate cell-to-cell signaling (1-3). While the structure and function of gap junctions and primary cilia is rather well understood, PD structure and function remain somewhat enigmatic.

PD were serendipitously discovered in the 1800s by the botanist Eduard Tangl while staining cell wall cellulose using organic pigments. To his surprise he observed cytoplasmic connections between plant cells in the endosperm (4). Similar structures were subsequently identified in many other plant species. Notably, analogous structures likely evolved independently at least six times, underlining their necessity for multicellularity (5). PDs connect neighboring cells by forming sophisticated channels lined by a contiguous plasma membrane. A special feature of higher plant PD is the desmotubule, a strand of apparently tightly appressed ER that extends through the central cavity (6-8). Since their discovery, plasmodesmata are considered as transport channels, enabling diffusion of small molecules below a certain size (9). They also seem to enable the passage of macromolecules, such as RNAs, peptides and proteins that are vital for plant development and physiological processes (10-12). Recently, tethering between the plasma membrane and the endoplasmic reticulum (ER) within PD was suggested as a mechanism to control PD pore size and Multiple-C2-domains-and-transmembrane-region-proteins (MCTP) were shown to play a major role in this process (13). However, the mechanism of transport through plasmodesmata is still unclear, as is their precise structural composition.

Proteomic analyses of PD enrichments provide a glimpse of the protein composition of these complex channels. Previous work identified PD components enriched from cell wall preparations (14-16). Some of these proposed PD proteins were also found enriched in detergent-resistant membrane (DRM) fractions. This may be due to the plasma membrane in PD being characterized by a distinct lipid composition (17, 18). PD proteomes were mainly defined by biochemical enrichment from cell wall preparation, for example in Arabidopsis (16, 19, 20), *Populus trichocarpa* (21), or *Nicotiana benthamiana* (22). Out of several thousand candidates identified in such proteomics analyses, only a small subset has in fact been validated with PD localization. A high-confidence definition of PD composition remains challenging due to the plasmodesmata being highly complex, comprising proteins of the cell wall, plasma membrane and endoplasmic reticulum. Biochemical preparations of PD obtained, for example, through cell wall enrichment or differential centrifugations contain a large proportion of co-purifying proteins from cellular domains other than PD. This requires quantitative methods distinguishing PD proteins from non-PD proteins. Comparative quantitative assessments of PD proteomes were performed, for example, by determining an enrichment ratio of proteins identified in PD compared to cell walls to define a ‘core PD proteome’ (13). Alternatively, a reduced abundance of PD proteins was observed in the loss of function mutant of CHOLINE TRANSPORTER 1 (*cher1-1*) and this was used to rank PD components (16). Another approach to characterize PD proteomes, explored protein recruitment to PD in response to viral infection (22). Viral movement between cells occurs through PD and thus proteins recruited or depleted from PD in response to viral infection were considered as candidate PD proteins. However, due to limited validation and limited number of species that have been used in PD studies, a reliable PD proteome with the probabilistic classification of core PD components and co-purifying proteins remains to be resolved.

Here, we used *Physcomitrium patens* as a model organism from the evolutionarily early land plants to establish an iterative discovery-verification workflow of the plasmodesmata proteome which uses experimental protein enrichment and a feature scoring algorithm in iteration with *in planta* localization. Our results give insights into the identified cell-wall modifying enzymes and allow conclusions on their roles during evolution of multicellularity.

## Results

### Detection of PD in *Physcomitrium*

To evaluate the presence of PD in *Physcomitrium*, transmission electron micrographs were generated from protonemata. Simple PD morphotypes were found at the interfaces between protonema cells at a relatively high density of 11.3 (± 1.4) PD µm^-2^ (Figure 1A, Supplementary Figure S2A) compared to PD densities in Arabidopsis roots, which ranged from 2.5 to 12.5 PD µm^-2^, depending on the cell-type interface (23). Protonemata PD were randomly distributed along the cell-to-cell interface and not clustered in pit fields (Supplementary Figure S2B). To identify PD protein candidates, a data-driven proteomics-based approach was used in iteration with systematic verification of PD localization.

**Figure 1:**
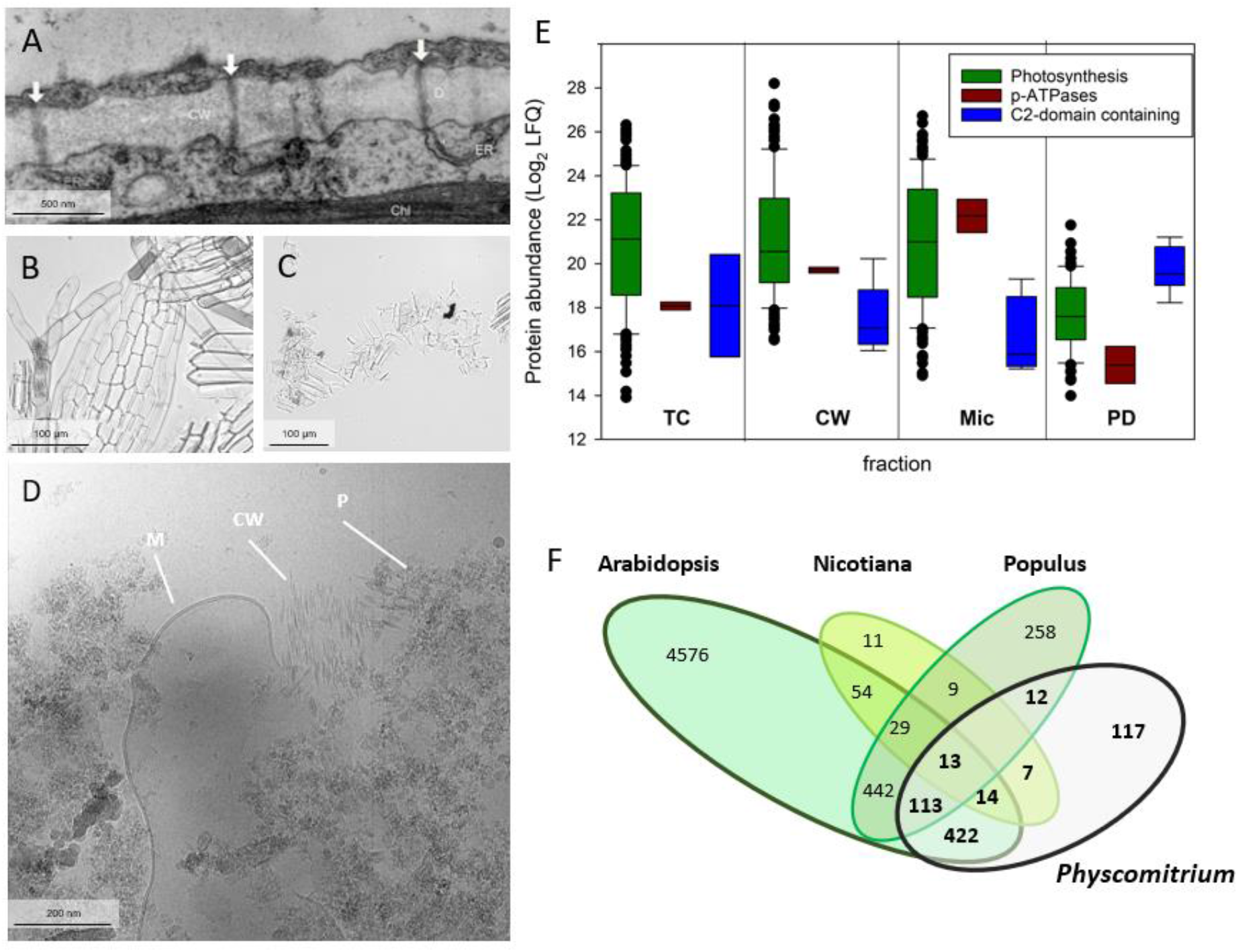
Proteins identified by the optimized protocol of PD enrichment from *Physcomitrium patens*. (**A**) Transmission electron micrograph of simple PD in *P. patens* protonemal cells. Arrows point three PD. CW, cell wall; ER, endoplasmic reticulum; Chl, chloroplast; Dt, desmotubule. Scale bar, 500 nm (**B**) Homogenized tissue (SI Materials and Methods). (**C**) Cell wall tissue after grinding in liquid nitrogen. (**D**) Electron micrograph of the PD pellet showing membranous components, protein aggregates and remaining cell wall (SI Materials and Methods). (**E**) Depletion of non-target proteins (e.g. photosynthetic proteins) and enrichment of target proteins (e.g. C2-domain containing proteins) in PD preparations. TC: total cell extract; CW: cell wall; Mic, microsomal membranes; PD: Plasmodesmata pellet. Scale bar, 200 nm. (**F**) Overlap of identified protein in PD pellets from *Physcomitrium* and published data sets from *Arabidopsis* (13, 16, 19, 20), *Populus trichocarpa* (21) and *Nicotiana benthamiana* (22). For Arabidopsis, the protein lists from four different studies were combined. For this comparison, Arabidopsis orthologs from respective species were used based on mappings available from Phytozome (70).

### Identification of PD protein candidates in *Physcomitrium*

Purification of PD remains challenging due to their small size, their incorporation in the cell wall and their dynamic proteomic composition. We achieved an enrichment of common PD proteins from *Physcomitrium* with concurrent depletion of typical contaminants. First step was disruption of plant tissue with a Potter Elvehjem Homogenizer in presence of high detergent concentrations. This gently broke the cells and removed organelles (16), while leaving membranes and cell walls largely intact (Figure 1B). Subsequent multiple rounds of grinding cell wall fractions in liquid nitrogen efficiently removed soluble protein contaminants and prepared cell walls for digestion by breaking remaining cell clusters (14). Washing with lower concentration of detergents prevented premature solubilization of PD membranes (14) before digestion of cell walls using driselase (Figure 1C). After ultracentrifugation, the resulting PD pellet contained small membranous components and protein aggregates with very little remains of cell walls (Figure 1D). Along the PD preparation workflow (Supplementary Figure S1) four protein fractions were collected as input for later PD Scoring: the first extract of cell wall proteins (CW), the total of microsomal membranes (Mic), the final pellet of PD enriched proteins (PD), and a total cell extract (TC). We observed a reduced abundance of photosynthetic proteins in PD pellets compared to TC or CW fractions, while proteins expected in PD, such as C2-domain containing proteins (13), increased in abundance (Figure 1E). Non-PD membrane proteins, such as plasma membrane ATPases, were most abundant in the microsomal fraction (Figure 1E).

Our PD preparation workflow identified 870 proteins in the final PD pellet. A comparison of published PD pellet proteomes from Arabidopsis (13, 16, 19, 20), *P. trichocarpa* (21), and *N. benthamiana* (22) revealed large overlap of 422 Arabidopsis orthologs in PD pellets of *Physcomitrium* and Arabidopsis (Figure 1F). Besides typical PD protein functions in cell wall modification or signaling, there was a high fraction (22%; n=38) of typical contaminants, such as ribosomes and photosynthetic proteins from light harvesting complexes and Calvin-Benson Cycle among orthologs found in at least three species. The fact that such non-target proteins made up a high percentage of the overlap in PD protein lists suggests that definition of PD proteins solely on the basis of discovery in multiple proteomics data sets lacks specificity and requires additional criteria for robust definition of PD composition.

### Definition of a high confidence PD proteome using a scoring system

To overcome the problem in distinguishing actual PD proteins from non-PD proteins in biochemical PD preparations, we developed a PD Score which is based on protein abundance data along the PD preparation workflow and on annotated features of a reference data set containing confirmed PD proteins from public sources.

We compiled the published proteome data sets in a relational database (PDDB http://pddb.uni-hohenheim.de, Supplementary Figure S3) and complemented 70 individual protein entries with literature evidence of experimental PD localization (Supplementary Table 1). These proteins were used as *bona fide* confirmed PD proteins and were further expanded by 115 proteins published as the PD core proteome (13) based on their high enrichment in PD fractions compared to cell wall or membrane fractions. Additional reliable PD proteins included 152 proteins with greater than two-fold depletion in the *cher1-1* mutant (16), and 56 proteins (52 unique Arabidopsis orthologs) with more than fivefold enrichment or depletion in PD in response to viral infection (22). The respective threshold values were chosen from the histogram of published quantitative values (Supplementary Table 1). The list of 201 proteins published as PD enriched from *P. trichocarpa* (21) was not included, since no enrichment ratios were published, making a quantitative re-evaluation difficult. Thus, the first PD scoring training data set comprised 338 unique Arabidopsis orthologs (Supplementary Table 1) coming from complementary data sources (Figure 2A).

**Figure 2:**
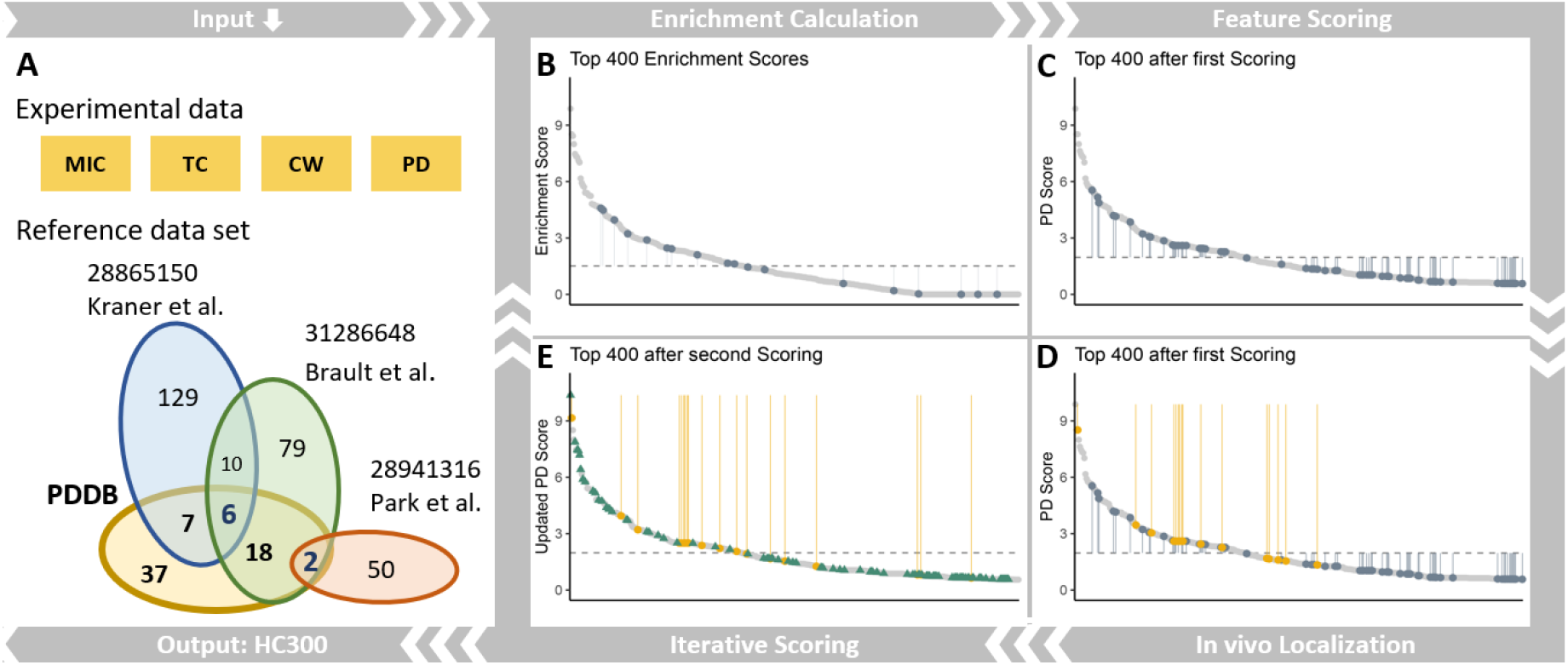
PD Scoring process visualized with the top 400 best scoring proteins. (**A**) Input data. consisting of protein abundances in MIC, TC, CW and PD fractions and the *bona fide* PD reference data set of PD proteins characterized by quantitative proteomics (13, 16, 22) (blue, green, red) and the additional PD localized proteins in PDDB from different literature sources (yellow). This data set was used as a PD positive set in the first round of PD scoring. (**B - E**) The components of the PD scoring process. The y-axis shows enrichment score or PD Score. Each of the top 400 scoring proteins is represented by a light grey dot. Proteins are ordered by score in descending order from left to right. Dark grey dots and vertical lines mark the scores of proteins from the reference data set. Yellow dots and vertical lines mark the scores of proteins that have been verified by *in vivo* localization. Grey dashed lines show mean scores. (**B**) Calculation of the enrichment score that is solely based on proteomics data. (**C**) PD Scores after including a first round of feature scoring. Higher values represent higher likelihood for PD presence. (**D**) Scores of newly PD localized proteins are highlighted before iterative scoring. (**E**) Iterative second PD Scoring including proteins experimentally localized to PD across the full range of PD candidate proteins. Green arrows indicate proteins which were not part of the reference data set but increased in score after iterative scoring.

The PD Score was calculated for each protein identified as the sum of an Enrichment Score and a Feature Score (Supplementary Figure S4). The Enrichment Score was derived from abundance of proteins identified in PD fractions compared to respective abundance in CW, MIC and TC. The feature score was calculated from frequencies of PFAM domains (http://www.pantherdb.org) and functional classifications by Mapman (24) in the reference data set of 338 *bona-fide* PD proteins. Consequently, proteins received higher PD Scores when they were enriched in the PD fraction and in addition shared features with reference PD proteins.

For 863 proteins found in the *Physcomitrium* PD pellet we were able to calculate an Enrichment Score and for 200 proteins the enrichment score was greater than 1 (Figure 2B). The full PD Score was calculated for 2053 out of the 4996 identified proteins (Supplementary Table 2). There were more proteins with a PD Score (Figure 2C) than with an Enrichment Score, because also proteins not quantified by an Enrichment Score were awarded a PD Score based on their features and can potentially be part of the PD proteome. Out of this PD score distribution, 147 proteins were cloned for validation of PD localization by transient expression in *N. benthamiana* (Figure 2D).

### Validation of PD localization using confocal microscopy

Putative PD-localized proteins were screened for colocalization with the callose-staining dye aniline blue as a marker of PD pit fields. Candidates were initially screened for PD localization and given a preliminary validation score (Supplementary Table 3). Proteins not colocalizing with aniline blue (112 candidates) were considered as non-PD proteins for the purpose of PD Scoring iteration. For the 36 proteins that colocalized with aniline blue, the degree of enrichment at PD compared to PM was calculated using our semi-automated PD indexing script. Proteins with a PD index > 1.1 (the highest mean score of the negative controls, Figure 3B) were considered enriched at PD compared to bulk PM. For comparison, aniline blue and the Arabidopsis protein PLASMODESMATA LOCATED PROTEIN 6 (PDLP6), a protein known to localize exclusively to PD (25), were used as positive controls. Aniline blue had a PD index of 4.4 while PDLP6 had a PD index of 2.5. Similarly, three proteins from the initial screen that did not show any obvious colocalization with aniline blue, localizing to either PM, vesicles or cytoplasm, were used as negative controls and had PD indices of approximately 1. Due to deficiencies in image quality, the PD index could not be calculated for all candidates deemed PD-localized by qualitative means. In total, we calculated PD indices for 30 of the 36 candidates that colocalized with aniline blue. For these candidates, PD indices ranged from 1.4 to 2.7. For the purposes of iterating the PD Scoring, all 36 candidates, including those not assigned a PD index, were treated as PD localized proteins.

**Figure 3:**
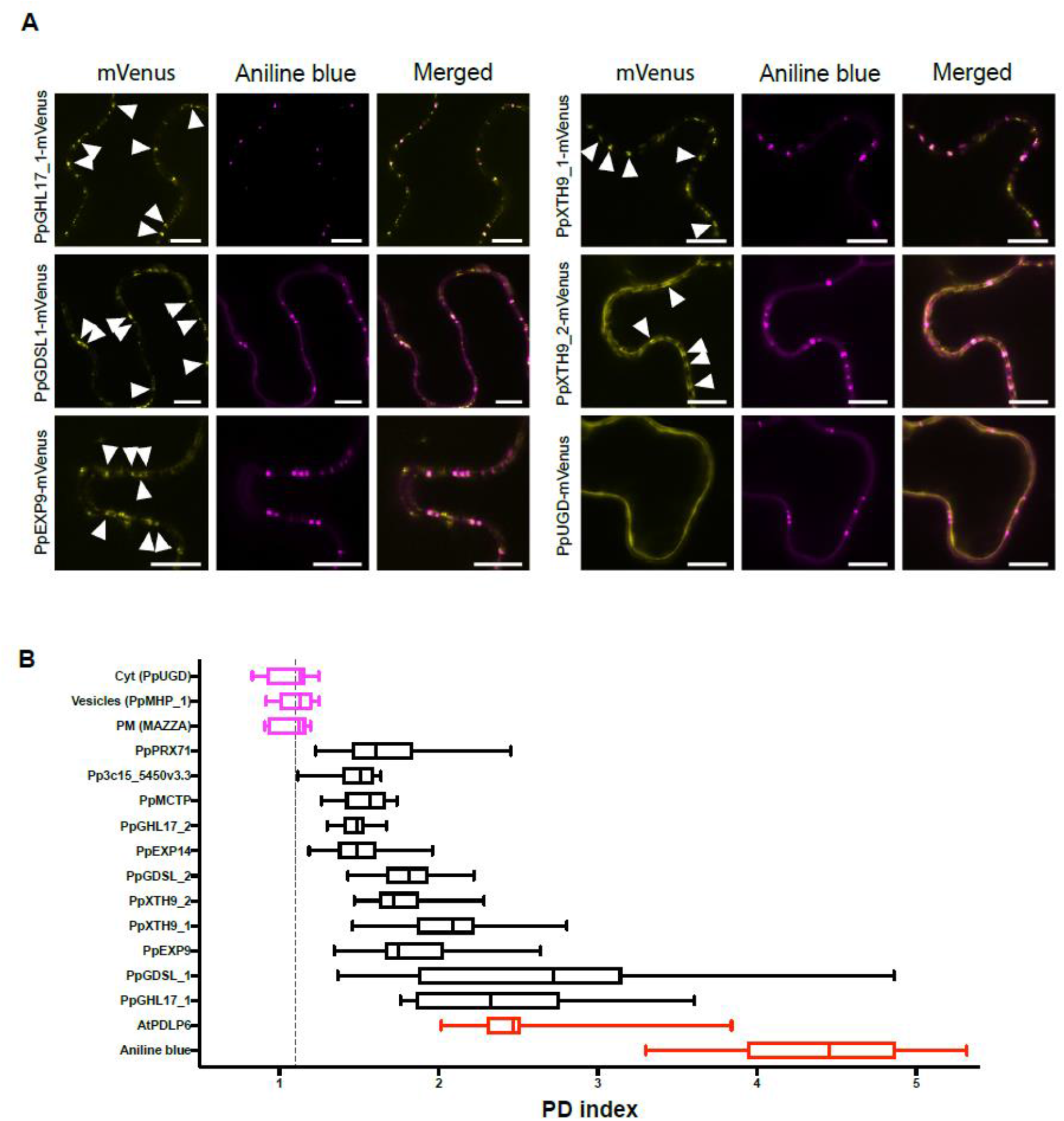
Selected candidate proteins from *P. patens* PD proteome located at plasmodesmata. Subcellular localization of PpGHL17_1 (Pp3c17_13760v3.2), PpGDSL1 (Pp3c2_2920v3.5), PpXTH_1 (Pp3c6_480v3.2), PpXTH9_2 (Pp3c14_17100v3.4), PpEXP9 (Pp3c18_1966v3.2) and PpUGD (Pp3c19_4880v3.3) in epidermal cells of *N. benthamiana* and observed at the confocal microscope. Proteins were C-terminally fused to mVenus and transiently expressed under a β-estradiol inducible promoter. (**A**) Single optical sections at cell-to-cell interface showing the co-localization of selected candidates with plasmodesmata marker aniline blue. All candidates showed an overlay with aniline blue (white arrowheads) except PpUGD which displayed cytosolic localization. Scale bars, 10 µm (**B**) Plasmodesmata index (PD index) of selected candidates, including those from panel A, showed an enrichment at PD similar to well-established plasmodesmata marker Arabidopsis PDLP6. PD markers aniline blue and PDLP6, and non-PD localized proteins Arabidopsis MAZZA (PM), PpMHP_1 (Pp3c16_460v3.7) (vesicles) and PpUGD (cytoplasm) are shown as red and blue box plots, respectively. In the box plot, median is denoted by a line and box and whiskers represent the 25th to 75th percentiles, and minimum to maximum distribution. Dashed line represents the threshold which a protein is considered PD-enriched and was set at 1.1 corresponding to the highest mean score of the non-PD localized proteins PpMHP_1.

### Refinement of a *Physcomitrium* high confidence PD proteome

The 36 newly confirmed *P. patens* PD proteins were used to improve PD scoring in an iterative second round. Among this set of proteins were 19 proteins which had not previously been considered as PD proteins in other species (Supplementary Table 4). For iterative scoring, the feature score component was modified by the contribution of additional PD localized proteins. As a result, most of the experimentally localized PD proteins increased in PD Score, as did other proteins with similar features (Figure 2E). This visualizes the key concept of PD scoring consisting of (i) protein abundance enrichment in PD fractions, (ii) identification of proteins with similar features as known reference proteins, and (iii) re-evaluating their PD Score based on features of experimentally validated PD-localized proteins.

The iterative PD Scores for proteins confirmed as PD localized in *N. benthamiana* clearly separated from PD Scores of protein which could not be localized to PD (Figure 4A). A threshold PD Score of 0.7 was chosen by carefully balancing the numbers of proteins not found with PD localization among the top-ranking proteins against proteins found with PD localization among lower ranking proteins. Among those 335 PD protein candidates with PD score greater than 0.7 (Supplementary Table 2) were 16 proteins which did not show PD localization in *N. benthamiana*. Although there may be many valid reasons for failed PD localization of these proteins, we termed these as false positives and they made up 3.9 % of the top 335 ranking PD protein subset. In turn, 28 PD localized proteins (1.6 %) were found among the 1718 proteins with a calculated PD Score less than 0.7, which were considered as false-negatives.

**Figure 4:**
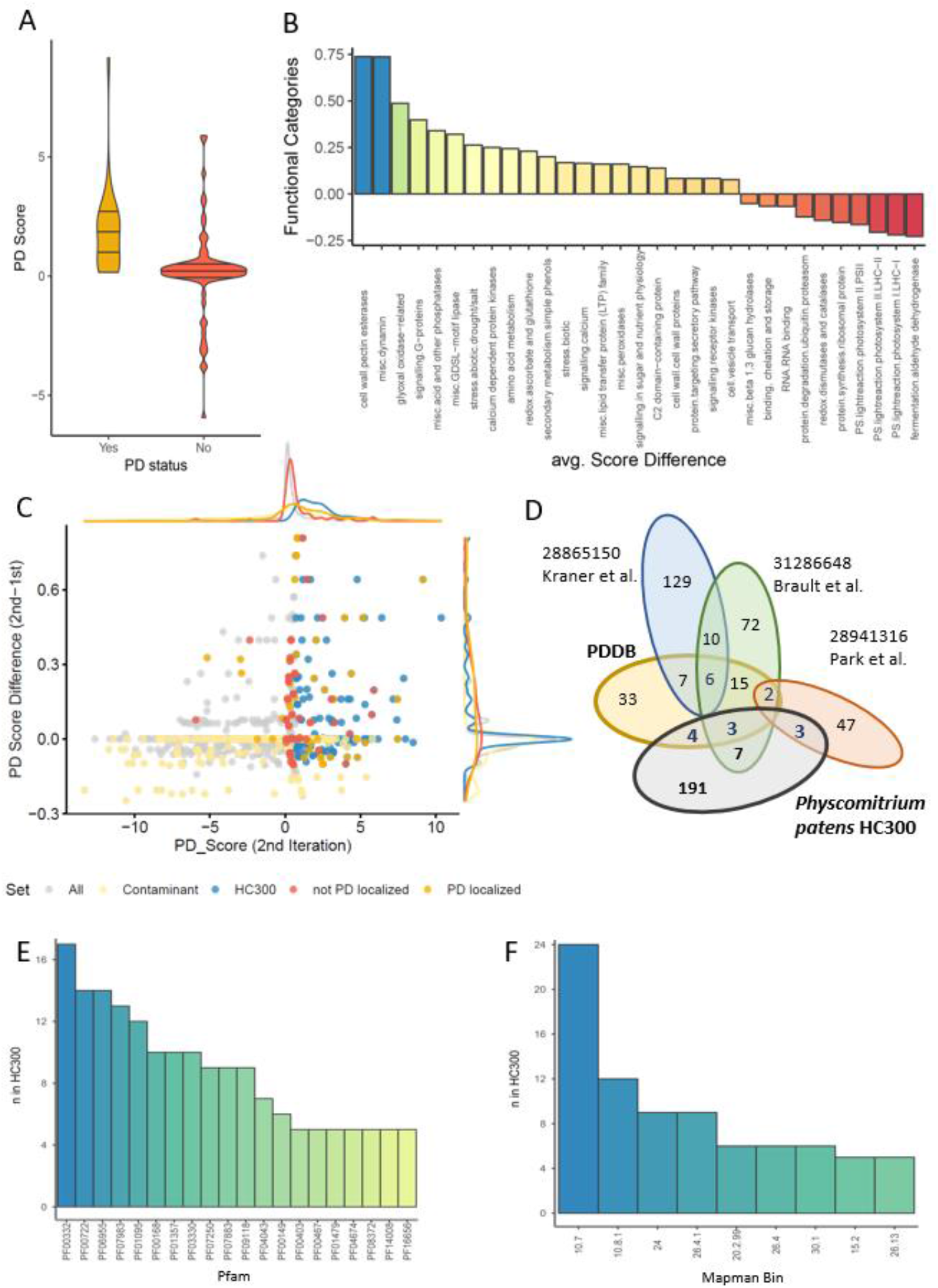
Characterization of *Physcomitrium* PD scored proteome. (**A**) violin plot of PD Scores (2^nd^ scoring) of proteins localized and non-localized to PD in *N. benthamiana*. (**B**) Averaged PD Score difference of iterative 2^nd^ scoring and 1^st^ scoring for proteins of different functional categories according to Mapman. (**C**) PD Score (>0.7) and PD Score difference (>-0.13) as criteria to define the high confidence *Physcomitrium* PD proteome. Blue: refined HC300 proteome. Red: photosynthetic proteins and ribosomal proteins as typical contaminants. Yellow: proteins with PD localization in *N. benthamiana*. (**D**) Venn diagram containing the refined high confidence PD proteome HC300 and the *bona fide* PD reference data set. (**E**) Mapman (24) functional categories significantly over-represented in HC300. 10.7, cell wall.modification; 10.8, cell wall.pectin esterases; 24, biodegradation of xenobiotics.glyoxal oxidases; 26.4, misc.beta 1,3 glucan hydrolases; 20.2 stress.abiotic; 30.1, signalling.in sugar and nutrient physiology; 15.2, metal handling.binding, chelation and storage; 26.13, misc.acid and other phosphatases. (**F**) PFAM domains (https://pfam.xfam.org) significantly over-represented in HC300. PF00332, Glyco_hydro_17; PF00722, Glyco_hydro_16; PF06955, XET_C; PF07983, X8; PF01095, Pectinesterase; PF00168, C2.

The 335 PD protein candidates were further refined by considering the PD Score difference between the second and the first round of PD Scoring, thus taking advantage of the iterative improvement of PD Scores after experimental validations. For protein functions expected among PD proteins, such as cell wall modifying proteins, signaling proteins, and C2-domain containing proteins, the PD Score improved throughout the iterative scoring process leading to high average PD Score differences (Figure 4B). In turn, for typical contaminant proteins, such as ribosomes, catalases, and photosynthetic proteins the PD Score was downgraded by the iterative process resulting in large negative PD Score differences (Figure 4B). By applying the PD Score difference as further refinement, we excluded proteins which had a strong negative PD Score difference (PD Score difference < -0.13) (Figure 4C). The resulting *Physcomitrium patens* High Confidence PD Proteome (HC300) contained 292 proteins, of which 273 proteins were annotated with Arabidopsis orthologs. Among the proteins in HC300, 39 proteins were direct orthologs to confirmed PD proteins from other species, and 191 proteins were not previously considered as high confidence PD proteins (Figure 4D). This additional refinement step efficiently removed typical contaminants, such as photosynthetic proteins and ribosomes. The content of contaminant proteins was found rather high (25%) among the proteins not considered as high confidence PD proteins while in HC300 the fraction of putative contaminants was reduced to 15%.

Within HC300, 62% of the proteins were found to have a signal peptide, 76% had at least one transmembrane domain, and 57% of the proteins were predicted to have intrinsic disordered regions (IDR). The PFAM X8 domain (PF07983), proposed with a role in protein targeting to PD through its signal sequences for a glycosylphosphatidylinositol (GPI) linkage, was found 14 times (5%) (Figure 4F) (26). Cell wall proteins (84 proteins; bins 10; p=5.5×10^−26^) were highly over-represented, as were glyoxal oxidases (bin 24; p=8.9×10^−13^) (Figure 4E). In agreement with the overall high content of cell wall related proteins, HC300 was significantly enriched with PFAM domains of glycosyl hydrolase 17 family proteins (PF00332, GHL17s), xyloglucan endotransglucosylases/hydrolases (PF06955, XTHs), and pectin esterase (PF1095). Moreover, four out of the 17 EXORDIUM/EXORDIUM-LIKE (EXO/EXL) proteins, whose Arabidopsis orthologs were previously shown to be located in the cell wall (27), were also found in HC300. Besides the cell wall proteins, HC300 contained five out of seven MCTP protein family members (Pp3c27_520V3.2.p, Pp3c14_25200V3.2.p, Pp3c10_11080V3.4.p, Pp3c16_9250V3.3.p, Pp3c27_540V3.2.p) as well as two other C2-domain containing proteins (Pp3c9_16970V3.3.p and Pp3c9_12510V3.5.p). PD localization of MCTP Pp3c10_11080V3.4.p was confirmed by transient expression assay (Supplementary Figure S5). In addition, three receptor kinases (Pp3c25_12800V3.2.p,Pp3c1_10970V3.4.p, Pp3c19_8660V3.4.p) were considered part of HC300 (Supplementary Table 2).

### Phylogenetic distribution of glycosyl hydrolase family 17

PD in land plants evolved to more complex structures compared to their counterparts found in *Characeae* algae (28). This might reflect adaptations to regulate PD conductance and the emergence of specialized metabolic enzymes. Deposition of callose at the neck region of PD is one of the major modes to gate PD (29). The maintenance of callose associated with PD is affected mainly by callose synthases and ß-1,3-glucanases. Among the candidates identified in HC300, there were 16 proteins annotated as ß-1,3-glucanases containing a glycosyl hydrolase family 17 (GHL17) domain (Supplementary Table 2). GHL17s are callose degrading enzymes located at the cell wall and associated to PD (30-32). Members of the GHL17 family were previously identified in PD proteome lists from Arabidopsis, *P. trichocarpa* and *N. benthamiana*, and 10 of them overlapped in at least two species. PpGHL17_16 (Pp3c16_16680V3.2.p) was the only member of the GHL17 family, for which orthologs were found in PD proteome lists of all four species. *Physcomitrium patens* HC300 contained three GHL17 orthologs (PpGHL17_2, PpGHL17_7, PpGHL17_9; Supplementary Table 5) which showed no overlap with PD proteomes from the higher angiosperms (Figure 5A).

**Figure 5:**
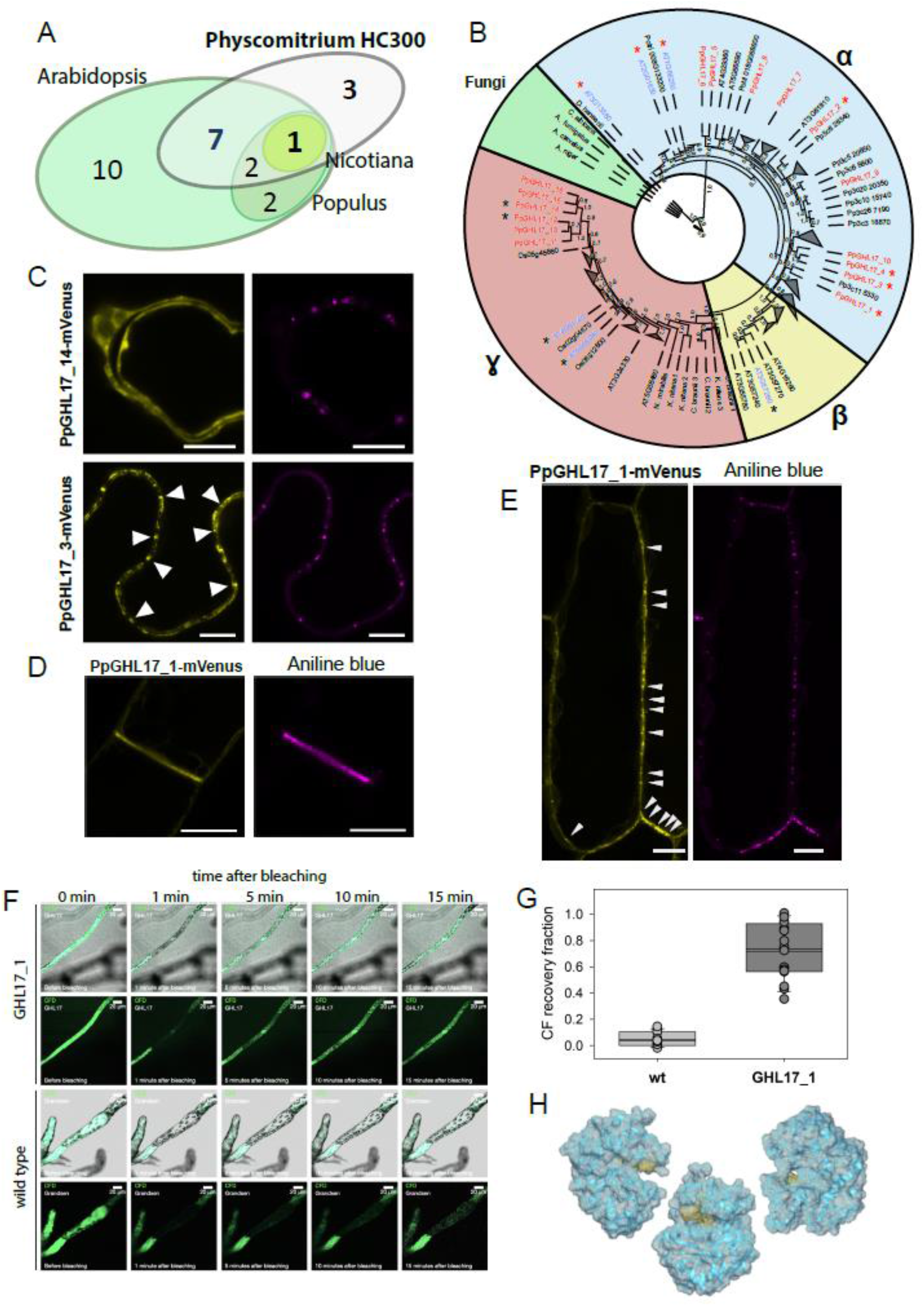
PD localization of GHL17 proteins correlates with phylogenetic distribution. (**A**) Members of the GHL17 family identified in PD proteomes from angiosperms (Arabidopsis, *Populus trichocarpa, Nicotiana benthamiana*) and the moss *Physcomitrium*. (**B**) Phylogenetic analysis of GHL17 sequences showed three major clades: α(yellow), β (blue) and *γ*(red), as defined by Gaudioso-Pedraza et al., 2014 (34). *P. patens* GHL17 proteins identified in the PD proteome are labeled in red and Arabidopsis orthologs previously assayed for localization colored in blue (30, 34). Red asterisks indicate *P. patens* and Arabidopsis PD-localized GHL17 proteins and group in clade α, whereas non-PD localized proteins are marked with black asterisks and cluster in either β or *γ*clades. Charophyta green algaes grouped in clade β. Fungi GHL17 sequences clustered separately (in green). The phylogenetic tree was generated with the maximum likelihood method implemented in PhyML (75). Support values from 1000 bootstrap samples are shown at the nodes. Clades which do not contain *P. patens* or Arabidopsis sequences were collapsed. (**C**) Subcellular localization in *N. benthamiana* epidermis cells of PpGHL17_14 (Pp3c6_17670v3.5) and PpGHL17_3 (Pp3c6_17670v3.5) mVenus fusion proteins belonging to β and αclades, respectively. PpGHL17_14 showed apoplasmic localization, whereas PpGHL17_3 displayed co-localization with aniline blue (pointed with white arrows). (**D**) Protonema cells of *P. patens* expressing PpGHL17_1-mVenus (Pp3c17_13760v3.2) showed an enrichment at the cell-to-cell interface. (**E**) Localization of PpGHL17_1-mVenus at PD pit fields (white arrowheads) in *P. patens* gametophore cells (SI Materials and Methods). Scale bars, 20 µm. (**F**) Images of CFDA/FRAP transport assays of *Physcomitrium* ecotype Grandsen wild type or over-expressing PpGHL17_1. (**G**) Quantification of CFDA transport rates in *P*.*patens* ecotype Grandsen wild type and over-expressing GHL17_1 presented as fraction of CF recovery in the photo-bleached protonema cells within 15 minutes (SI Materials and Methods). Values represent the mean ± SE of 16 experiments for each line. (**H**) AlphaFold model of *Pp*GHL17 αclade glycoside hydrolase domains (cyan) with the conserved IFALFNE(N) motif (yellow).

Based on sequence similarity, GHL17 proteins of representative species from charophytic algae and embryophytes, including *P. patens*, were classified into three major clades (33, 34). To get insights into their role in evolution of multicellularity, a new phylogenetic analysis of the GHL17 family was performed including *P. patens*, Arabidopsis, *P. trichocarpa, O. sativa* and representative species from green algae (Figure 5B, Supplementary Figure S6A). Unexpectedly, no GHL17 orthologs could be identified in the unicellular algae *Mesostigma viride* and *Chlamydomonas reinhardtii*, which both are ancestral species in *Streptophyta* and *Chlorophyta* clades. This suggests that the GHL17 family emerged in multicellular organisms.

GHL17s from charophytic algae (*K. nitens, C. braunii*, and *N. mirabilis*) were present only near the base of the gamma clade, likely representing the ancestral type of GHL17 proteins (Figure 5B). The alpha clade only contained GHL17s from embryophytes, suggesting that this group appeared during land colonization. Three of the Arabidopsis GHL17 proteins from the alpha clade were previously described to localize at PD, suggesting that phylogenetic distribution might correlate with the subcellular localization either at PD or apoplasm (32, 35). To test this hypothesis, we studied the subcellular localization of six *P. patens* GHL17 representatives, four from the alpha clade and two from the gamma clade by transient expression in *N. benthamiana*. Transient expression of gamma clustered PpGHL17_14-mVenus and PpGHL17_12-mVenus showed apoplasmic localization (Figure 5C, Supplementary Figure S6B), whereas PpGHL17_1-mVenus, PpGHL17_2-mVenus, PpGHL17_3-mVenus and PpGHL17_4-mVenus from the alpha clade were found with punctuated pattern at the cell periphery and overlaid with aniline blue (Figure 5C, Supplementary Figure S6B). Since transient expression assays can lead to mis-localization, and targeting mechanisms to PD might not be conserved between *N. benthamiana* and *P. patens*, we obtained a *P. patens* stable transgenic line expressing UBQ:PpGHL17_1-mVenus to further validate PD localization. The subcellular localization of PpGHL17_1-mVenus in the stable line confirmed PD localization found in transient expression assays (Figure 5D, E). PpGHL17_1-mVenus localized at the cell-to-cell interface in protonema cells and co-localized with the PD marker in gametophore cells. In addition, when over-expressing PD-localized PpGHL17_1 in *Physcomitrium*, diffusion of the PD mobile dye carboxyfluorescein diacetate (CFDA) increased compared to wild type ecotype (Figure 5 F,G) supporting a role of GHL17s in PD opening through the degradation of callose.

### Xyloglucan endotransglucosylases/hydrolases

XTHs are enzymes involved in xyloglucan metabolism, important for the control of cell wall strength and extensibility during growth and development. In Arabidopsis, XTHs were shown to be upregulated in the *ise2* mutants, which displayed an increase of cell-to-cell transport of fluorescent probes (36). We found 12 members of the XTH family in HC300, six of which were confirmed as PD-localized (Figure 3A, Supplementary Figure S7A). The molecular phylogeny of XTH genes was previously shown to segregate in three major groups (I, II and III), and an isolated small group named “ancestral” (37-39). All *Physcomitrium* XTH genes, except Pp3c13_11910 and Pp3c3_9730, clustered in the group I. The PpXTHs identified in HC300 were exclusively from group I (Supplementary Figure S7B), indicating a correlation between PD localization and phylogenetic distribution to group I.

### EXORDIUM and EXORDIUM-Like proteins

Four members of the Arabidopsis Exordium (EXO) and Exordium-Like (EXL) protein family were identified in HC300: Pp3c19_8770V3.5 (PpEXO_1), Pp3c10_8680V3.4 and Pp3c14_6120V3.3 (AtEXO ortholog by protein BLAST), and Pp3c22_12550V3.1 (AtEXL2 by protein BLAST). EXL2 was previously found in Arabidopsis and *P. trichocarpa* PD proteomes (20, 21). Interestingly, several members of this family were also found in Arabidopsis cell wall proteomes (AtEXO, AtEXL1, AtEXL2, AtEXL3, AtEXO4) and membrane proteome (AtEXL4) (19, 40-42). Here, PpEXO_1 and Pp3c22_12550V3.1 were found among the top 10 PD Score ranks in HC300 (Supplementary Table 2), and was localized at PD in transient expression assays in *N. benthamiana* (Supplementary Figure S8).

To understand functions of this protein family in PD, we looked for EXO/EXL orthologs within the kingdoms of life. No member of the EXO/EXL family was identified among the bacteria, protists (including Chromista and Ochrophyta), fungi and *animalia* kingdoms. In the *plantae* kingdom, EXO orthologs were found in all investigated embryophytes, from bryophytes to angiosperms. However, EXO members were scarce in algae. We could not identify orthologs in unicellular (*Cyanidioschyzon merolae*) and multicellular (*Chondrus crispus*) Rhodophyta. Among the 15 Chlorophyta species investigated, EXO orthologs were found only in the Trebouxiophyceae *Coccomyxa subellipsoidea*, despite it being a unicellular algae. Finally, most of non-embryophyte EXO proteins were found in charophytic algae, which together with the embryophytes form the Streptophyta lineage. Among the Charophyta, EXO proteins were identified in the non-PD-containing species *K. nitens*, in the PD-containing species *C. braunii*, as well as in the unicellular species *Spirogloea muscicola*, which is thought to be the closest sister species of the embryophytes. However, no ortholog could be found in *Mesostigma viride* and *Chlorokybus atmophyticus* which diverged earlier within the Charophyta lineage. EXO proteins thus appeared before the emergence of PD, but the presence of orthologs in only one of the investigated Chlorophyta species and their absence in early diverging Charaphytes remains puzzling.

A phylogenetic analysis was undertaken to study sequence divergence along the EXO family evolution. EXO proteins from representative species of the embryophytes (17 from *P. patens*, 8 from Arabidopsis, 12 from *O. sativa*, 17 from *P. trichocarpa*) as well as all EXO proteins identified in algae were therefore included in this analysis (Supplementary Figure S8). We found that EXOs are distributed within three clades. While clade I contained only EXO sequences from algae including both Chlorophyta and Charaphyta, clade II contained EXO sequences from all the investigated species, distributed between two subclades. Subclade II-a exclusively included EXOs from the algae *S. muscicola*, and subclade II-b contained embryophyte sequences. Such clustering suggests that embryophyte EXOs found in clade II were closer to the ancestral EXOs than embryophyte EXOs found in clade III, which contained EXO sequences from all investigated embryophyte species. The four *P. patens* EXO proteins identified in HC300 were all found in a subclade of clade III which seems to correspond to a cluster of EXO orthologs from seedless plants previously defined by the EXO sequences of *P. patens* and *Selaginella moellendorffii* (43). Interestingly, also EXOs from Arabidopsis and *P. trichocarpa* previously identified in PD proteomes, were found out of clade III (Supplementary Figure S8).

## Discussion

### PD scoring algorithm to define PD proteomes

Plasmodesmata are nanometer-small structures of high structural complexity. They contain different types of cellular membranes and protein composition depends on environmental cues, tissue, and/or developmental stage. Thus, we require approaches which tag likely PD localized proteins and separate them from likely non-PD proteins. The mere identification of a protein in biochemically enriched PD preparations is not sufficient to conclude its PD localization without additional information. We present a scoring method consisting of a biochemical component based on protein enrichment in PD preparations and a bioinformatic component based on protein features. We used a curated set of *bona fide* PD proteins to identify overrepresented protein features used for scoring. In this way, scoring can be iteratively improved by improved information in the input data set of known PD proteins. We demonstrated the improvement of the scoring algorithm after extensive protein localization and used this information to refine a high confidence PD proteome of *Physcomitrium* (HC300). A previous approach (44) also applied a bioinformatic strategy to predict PD proteins based on protein features found in a subset of the published proteomes (13, 20, 22). However, the authors did not consider biochemical aspects (i.e., enrichment in biochemical PD fractions) and they did not benchmark the predictions against experimental protein localization.

No biochemical method yields pure PD. Thus, highly abundant proteins such as photosynthetic proteins or ribosomes will always be detected in PD preparations. Upon stepwise refinement of the *Physcomitrium* HC300 PD proteome based on PD Score and PD Score Difference, the overlap with other quantitative PD proteomes (16, 22) became smaller at each refinement step (Figure 4D, Supplementary Figure S9). This supports our finding that the overlap between unweighted PD proteome lists contains a high proportion of such contaminant proteins. The highest overlap of our *Physcomitrium* HC300 remained with the published core proteome of Arabidopsis as concluded from enrichment of proteins in PD preparation (13), and enrichment values were a component also of the PD scoring applied here.

### Large scale PD localization in N.benthamiana

Despite improvement in distinguishing likely PD proteins from likely non-PD proteins by the PD Scoring, experimental localization remains important. For example, we tested five members of the GHL17 family of glycosyl hydrolases for PD localization with positive results for three proteins and no PD localization in *N. benthamiana* for two family members. Overall, the transient localization of *Physcomitrium* proteins in the *N. benthamiana* system proved efficient and applicable also on a larger scale, and, most importantly, showed that targeting mechanisms to PD are conserved between early land plants and vascular plants. However, we cannot exclude that false negatives occurred, due to the absence of required chaperones or mis-interpretation of sorting signals in this heterologous system. In turn, false positives could have resulted from over-expression or mis-localization. To address such caveats, we used an inducible promoter and appropriate induction times to minimize overexpression artifacts. Fluorescent protein fusions at different positions may affect localization and the proteins may also function at more than one subcellular localization, as was already shown for MCTPs (45). Nevertheless, of the 147 proteins tested, the majority (75%) did not colocalize with aniline blue and were thus considered as non-PD-localized. Given that PD are extremely small structures and their biochemical preparation remains challenging, the fraction of about 25% of PD candidates from biochemical preparations being experimentally PD localized in *N. benthamiana* seems reasonable.

### Protein composition of the P. patens PD

Our high confidence subset of the *P. patens* PD proteome contains a high number (n=84) of cell wall and cell wall modifying proteins. Among these, the largest group comprised glycosyl hydrolases from the GHL17 subfamily. The GHL17 family is an excellent example of a large protein family which throughout evolution diversified in its subcellular localization, with one clade becoming PD localized. This subfamily can also be taken as an example highlighting the two components of the PD Scoring, the biochemical enrichment in PD, and a feature scoring component. The feature scoring component alone would not have been able to differentiate proteins from the two clades of GHL17 proteins. However, the whole PD Score revealed an important means to differentiate PD localized and non-PD-localized GHL17s. Predictions of structural properties suggest that most of the GHL17s of the beta clade have an X8 domain and are likely to be GPI anchored. GPI anchors were previously suggested to be a PD sorting signal (46). The alpha clade of PpGHL17s appeared to be more diverse, with only some family members being predicted to have GPI attachment signals and X8 domains (Supplementary Table 5). In search of a potential PD localization signal, we modeled structures of alpha-clade PpGHL17s (Figure 5H). The sequence motif IFALFENE(N) was over-represented among this group of GHL17s and was absent in the members of the gamma clade. However, this motif was predicted to be only partially surface-exposed as part of the 40 Å cleft of the glycosyl hydrolase domain and is therefore more likely to be related to enzyme function rather than localization (Figure 5H).

In the HC300 PD proteome we identified four out of the 17 *P. patens* EXO/EXL proteins. The ortholog to Pp3c18_22500V3.3.p is named PHOSPHATE INDUCED 1 (PHI-1) and was initially identified in tobacco BY-2 cell suspension cultures among genes induced after recovery from phosphate-induced starvation (47). In Arabidopsis, EXO/EXLs were identified as potential mediators of brassinosteroid-promoted growth (48, 49). Indeed, *AtEXO* and *AtEXL1* expression is induced by brassinosteroids, and *AtEXO* regulates other brassinosteroid-responsive genes (48, 49). While overexpression of *AtEXO* promotes shoot and root growth, its loss of function lead to a reduction of growth and biomass production resulting from a defect in cell expansion (27). The expression of several *AtEXLs* is also controlled by carbon metabolism and energy status, and is required during adaptation to carbon and energy-limiting growth conditions (48). Although the physiological functions of EXO proteins are still unclear, it was also proposed that AtEXO could connect extracellular carbon status to growth responses (27, 48, 50).

Recently, bud dormancy was shown to be associated with a molecular carbon starvation response which includes an increase in *EXL2* and *EXL4* RNA levels (51). During bud dormancy, meristematic cells are symplasmically isolated through callose-dependent PD obstruction (52-55). Photoperiod and gibberellic acid mediate opening of PD by inducing the expression of GHL17s. GHL17s then promote callose hydrolysis at PD and thereby contribute to dormancy release (56). Because of the increasing evidence of EXO/EXL localization at PD, the involvement of these proteins in sensing of carbon status and regulation of growth response might need to be revisited in the context of symplasmic communication. Indeed, the mechanisms by which these proteins mediate their physiological functions remain to be elucidated. Contrary to what has been previously reported (43), we also found EXO/EXL orthologs in algae. However, EXO/EXL identified in the PD proteome do not cluster with the algae members, suggesting that EXO/EXL PD localization might have emerged later during evolution of embryophytes. The EXLs are not predicted to have GPI anchors, but they contain an N-terminal alpha helix which is predicted to contain an ER/secretory pathway targeting signal.

Here, we used strict scoring thresholds to define the high confidence set of proteins likely present in PD of *Physcomitrium patens*. However, the drawback of stringent thresholding lies in accepting potential losses of loosely associated proteins or proteins dynamically associated with PD. Similar effects were discussed previously for interactome data: proteins identified with high reproducibility and high confidence tended to make up stable protein complexes, such as ribosomes, respiratory chain or photosynthetic complexes (57). In turn, proteins with higher variability among replicates and lower confidence values tended to contain more protein functions for which dynamic or conditional interactions are expected, such as for example signaling proteins (57). A tradeoff between accuracy and coverage has also been discussed for yeast interactome data (58). In analogy, HC300 is rich in cell wall proteins and structural elements (e.g. the C2-domain containing proteins) but contains a rather low fraction of signaling elements such as receptor kinases. We found only one member of the GDSL-motif lipase/hydrolase family proteins in HC300, while 9 other identified family members were not considered part of the high confidence proteome based on their PD Score rank, although two of them were experimentally localized to PD. Similarly, also dynamins, RAN-GTPases, and glyoxal oxidase-related proteins were not classified among the HC300 after PD Score ranking (Supplementary Table 2). However, members of these protein families also showed a PD Score Difference greater than 0.3, indicating highly improved ranking based on features of experimental localizations. We therefore considered these protein families part of an extended PD proteome (Supplementary Table 2).

### Evolution of PD localization within large protein families

Symplasmic transport is highly regulated and involved in numerous processes in plant development, for example, bud dormancy and meristem activity. Thus, some of the PD proteins are involved in transport of signaling molecules and resource sharing and/or the regulation thereof. These functions must have evolved and diversified during evolution of multicellularity. The *Physcomitrium patens* HC300 contains several members of larger protein families, exemplified by GHL17s, XTHs and EXO/EXLs, each of which diverged into several phylogenetic clades. Interestingly, in all cases, PD localized members grouped in a distinct clade, often – as in the GHL17s – separating PD localized and non-PD localized family members. We hypothesize that a co-evolution of PD functions and the respective protein localization took place, leading to diversification in the respective protein families.

GHL17s and XTHs are considered important in controlling PD gating by affecting callose formation. GHL17s as callose degrading enzymes are considered to be involved in keeping PD in an open state (30-32)(Figure 5F,G), while for XTHs a role in cell wall remodeling during formation of secondary PD was proposed (59). Similarly, a callose synthase ortholog has recently been characterized with critical functions in enabling sucrose transport in rice (60). The presence of large numbers of proteins involved in cell wall synthesis and remodeling in the *Physcomitrium* PD proteome may reflect the need to dynamically modify the composition of the PD associated cell wall regions independently of other regions.

## Conclusion

Our work provides insights into the composition of PD in the bryophyte *P. patens* based on an iterative workflow of biochemical enrichment, feature scoring and verification of PD location by fluorescent microscopy. To enable PD feature scoring by use of public data sets of PD proteomes from other plants, we have established a public database, PDDB (http://pddb.uni-hohenheim.de), in which *bona fide* confirmed PD proteins are separately flagged. Our high confidence PD proteome HC300 further provides insights into large families of cell wall proteins which have diversified into PD-localized and non-PD-localized members throughout the evolution of multicellularity.

## Materials and Methods

### Plant material and culture conditions

*Physcomitrium patens* ecotype Grandsen cultures were obtained from the International Moss Stock Center (IMSC; Freiburg). *P. patens* was cultured on modified BCD medium containing 5 mM diammonium tartrate (61) (SI Materials and Methods). *N. benthamiana* seeds were soaked in water for 24 h at 20°C and then grown on peat moss substrate in a greenhouse for 3 weeks under long day conditions (SI Materials and Methods).

### Protein isolation

Total protein was extracted in a urea thiourea environment (62). Microsomal fraction isolation followed established protocols (63). Cell wall extracts were obtained after cell homogenization with a Potter-Elvehjem PTFE tissue grinder in high detergent environment (16) and centrifugation at 1000 x g. The cell wall pellet was subsequently ground in liquid nitrogen (14). Cell wall fragments were enzymatically digested with driselase and the released PD membranes were pelleted by ultracentrifugation at 110,000 x g (SI Materials and Methods).

### LC-MS/MS analysis of peptides

Proteins were trypsin-digested and desalted (64) and peptide mixtures were analyzed on a nanoflow UPLC-coupled Orbitrap hybrid mass spectrometer (Q-exactive HF, Thermo Scientific) (SI Materials and Methods). Spectra were matched against the *Physcomitrium patens* proteome (Ppatens_318_v3.3.protein.fasta, 87533 entries) using Maxquant MaxQuant version 2.0.3.0 (65). Mass spectrometry proteomics data were deposited to the ProteomeXchange Consortium via the PRIDE partner repository (66) with the dataset identifier PXD032820.

### PD Score

The PD Score is the sum of two components, the enrichment score (Eq.1) and the feature score (Eq.2 and 3). LFQ values from the TC, MIC, CW and PD fractions derived from MaxQuant (67) were log_2_-transformed before further calculations. Fraction means were calculated and normalized using z-Score (Supplementary Figure S4).

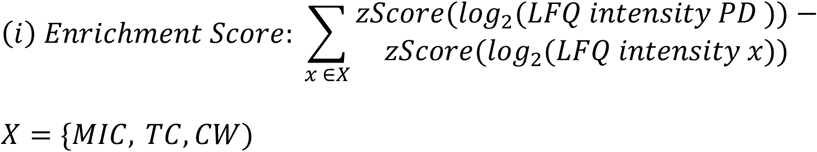

Feature scores were based on the frequency of protein functions and PFAM domains (https://pfam.xfam.org/) found in the reference data set and validated PD candidates. The feature score is the sum of feature frequencies of PD protein associated features found in the scored protein.

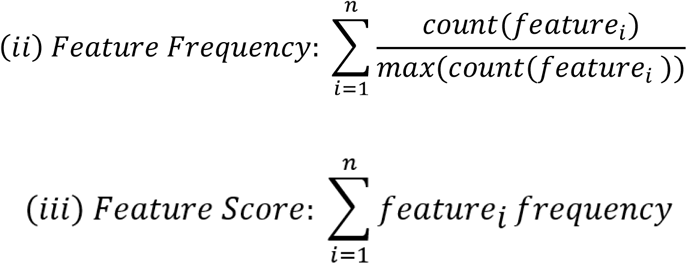

### Quantification of PD candidate localization

Confocal images were screened visually for colocalization of the protein of interest and aniline blue. Candidates showing colocalization with aniline blue were quantitatively scored by the so-called PD index (68). To minimize bias in this quantification, we used a semi-automated approach of an in-house-generated macro for the *Fiji* software package (69) (SI Materials and Methods).

### Phylogenetic analysis

GHL17s, XTHs and EXOs protein sequences were retrieved from Phytozome, Ensembl, Phycocosm and http://genome.microbedb.jp/klebsormidium databases (70-73). All protein sequences were subjected to protein domain prediction at InterPro server (74). Maximum likelihood trees were generated using the NGPhylogeny.fr tool (75) and visualized and edited in iTOL (76).

## Supporting information

Supplementary Figure S1_PD prep workflow

Supplementary Figure S2_TEM

Supplementary Figure S3_PDDB

Supplementary Figure S4_PD scoring formeln

Supplementary Figure S5_more localizations

Supplementary Figure S6_GHL17s full

Supplementary Figure S7_XTHs

Supplementary Figure S8_EXO

Supplementary Figure S9_venn diagrams

Supplementary Table 1_published data sets

Supplementary Table 3_PD validation scale

Supplementary Table 4_verified proteins

Supplementary Table 5_GHLs, XTHs, EXOs

Supplementary Table 6_primers

Supplementary Table 2_PD scoring list

## Acknowledgements

We thank Jan Weber for excellent assistance during protein preparations of PD fractions, Ronja Lange for excellent assistance in cloning of *Physcomitrium* genes. pTH Ubi-Gate plasmid was a kind gift from Prof. Bezanilla.

This work received funding from the European Research Council (ERC) and the Marie Skłodowska-Curie under the European Union’s Horizon 2020 research and innovation programme (Grant agreements “SymPore” No. 951292 to WBF, RS, WS, WB, “PDgate” No. 101023981 to MM, and “CLAVHUB” No. 101023589 to VH, respectively). This research was supported by the Deutsche Forschungsgemeinschaft (DFG, German Research Foundation) under Germany’s Excellence Strategy – EXC-2048/1 – project ID 390686111 (RS, SH and WF), and the Alexander von Humboldt Professorship (WBF).

## Notes

### Competing Interest Statement

The authors have declared no competing interest.

