## Supplementary material for "A high confidence *Physcomitrium patens* plasmodesmata proteome by iterative scoring and validation reveals diversification of cell wall proteins during evolution": SI Appendix

**Supplementary Table 1 "published data sets"**: The set of 338 proteins considered as *bona fide* PD proteins in the first round of PD relevance scoring.

**Supplementary Table 2 "PD scoring list"**: *Physcomitrium* proteins for which a PD Score was calculated and refined high confidence PD proteome HC300.

**Supplementary Table 3 "PD validation scale"**: PD localization was estimated based on a 0 to 10 scale by co-localizing the PD candidate and the PD marker aniline blue in one cell. The scores were as follows: 0 indicated none ROI of the PD candidate transiently expressed in *N. benthamiana* overlays with aniline blue; 1 indicates less than 3 ROIs overlaying with aniline blue; and so forth until a score of 9 which indicates more than 10 ROIs overlapping with the PD marker. Only those cells with all the ROIs overlaying aniline blue scored 10.

**Supplementary Table 4 "verified proteins":** The list of *Physcomitrium patens* proteins tested for PD localization.

**Supplementary Table 5 "GHLs, XTHs, EXOs":** Information on the glycosyl hydrolase family 17. List of proteins with a GHL17 PFAM domain (PF00332) identified in the Physcomitrium HC300 and in other species PD proteome lists, and list of GHL17, XTH, and EXO sequences used for constructing the phylogenetic trees.

**Supplementary Table 6 "primers"**: primers used for cloning of *P.patens* PD candidate genes.

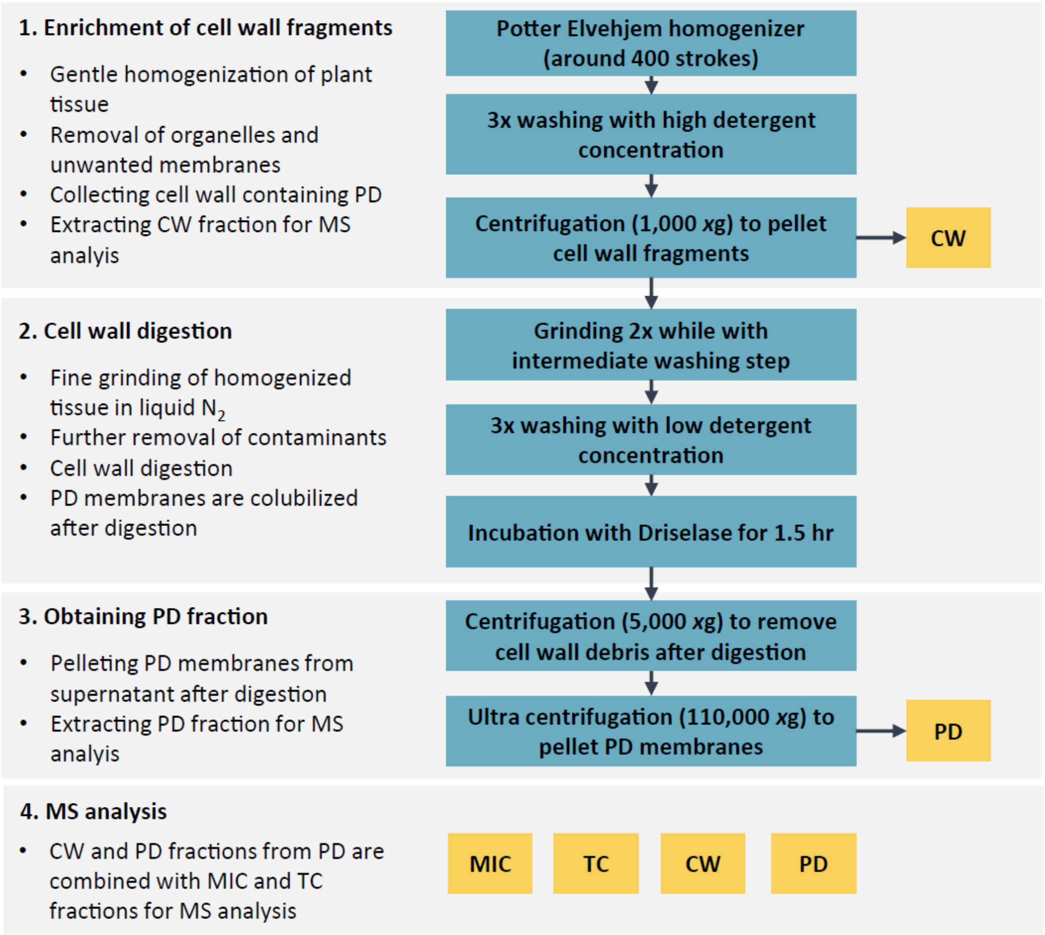


**Supplementary Figure S1**: Enrichment protocol of PD from *Physcomitrium patens* highlighting key features of the protocol and sampling stages along with the PD enrichment.

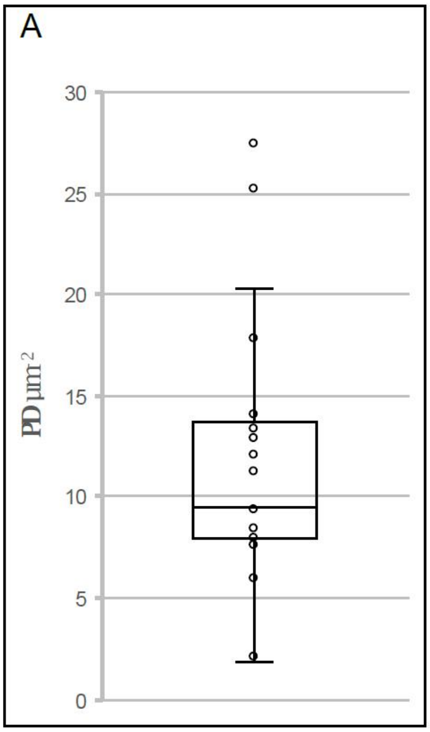


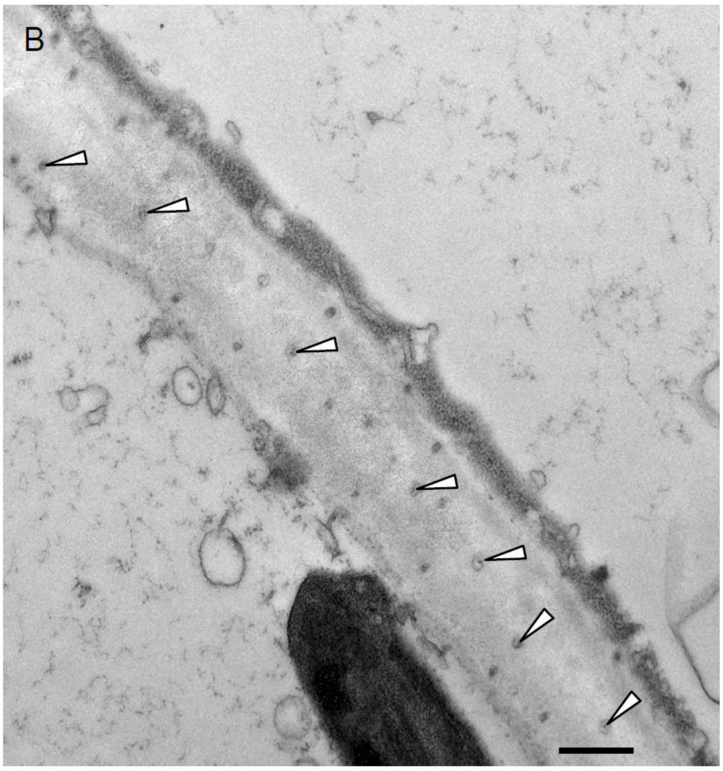


**Supplementary. Figure S2:** (**A**) PD density in protonema cell-to-cell interface as determined by TEM analysis. The box and whisker represent the 25 to 75 percentile and minimum-maximum distributions of the data, respectively. Median is indicated by line and individual values as empty circles. (**B**) Cross-section through simple PD between two protonema cells. Desmotubule (pointed with arrowheads) is visible in the central part of the lumen as a higher electron density. Scale bar, 500 nm.

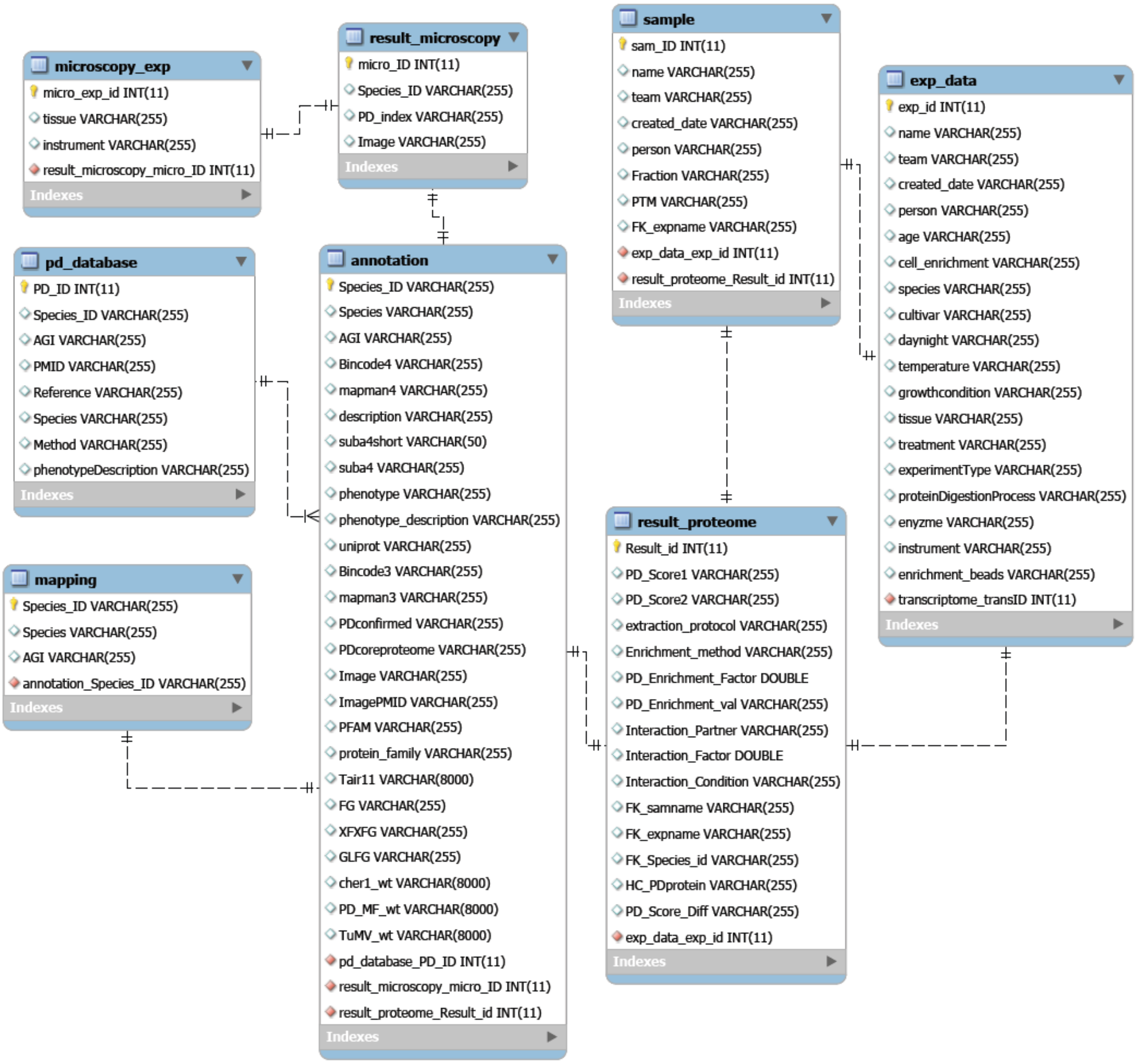


**Supplementary Figure S3**: Structure of the core PDDB database.

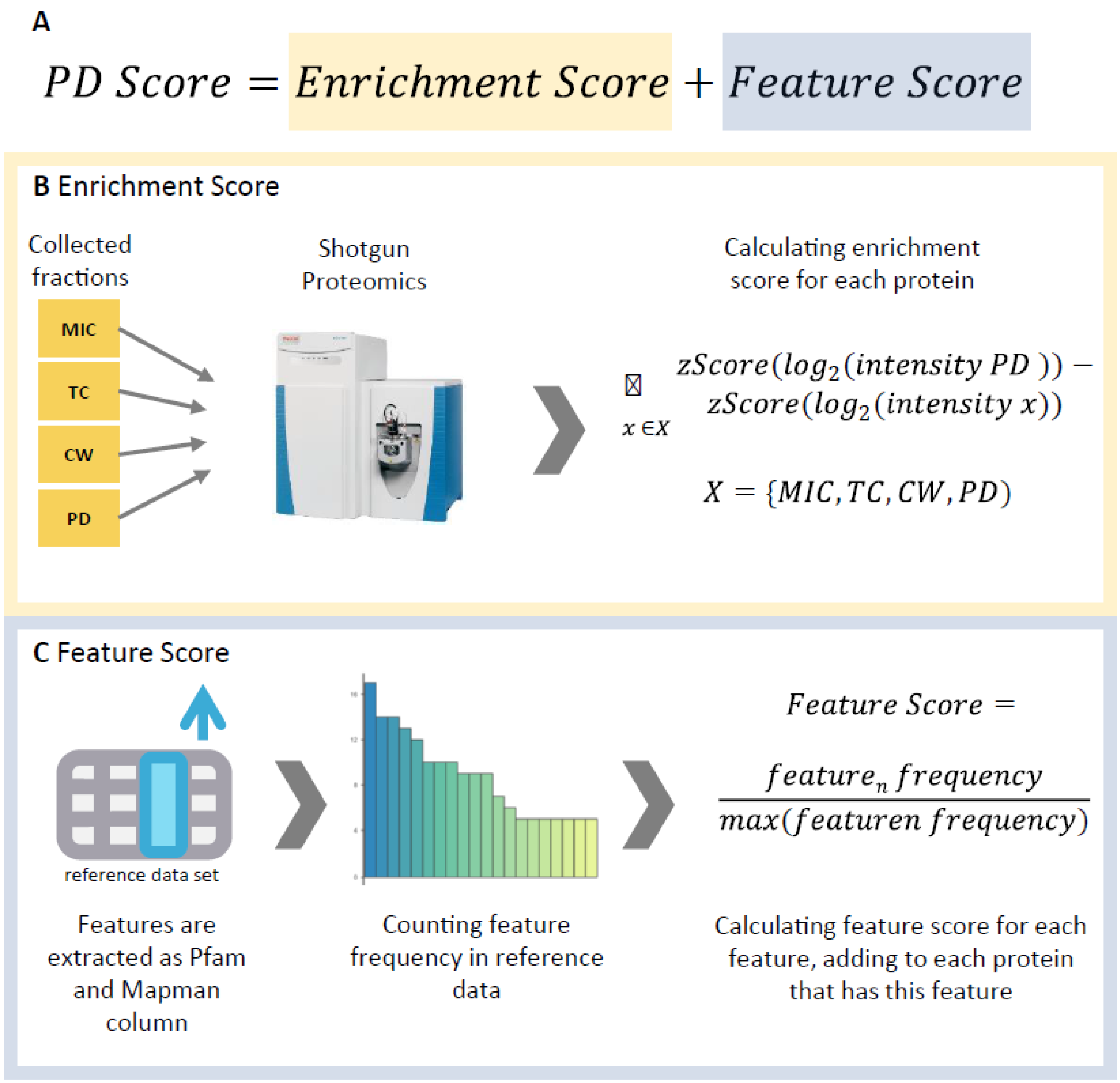


**Supplementary Figure S4**: Details on the PD Score. (**A**) calculation of the enrichment score. (**B**) calculation of the feature score.

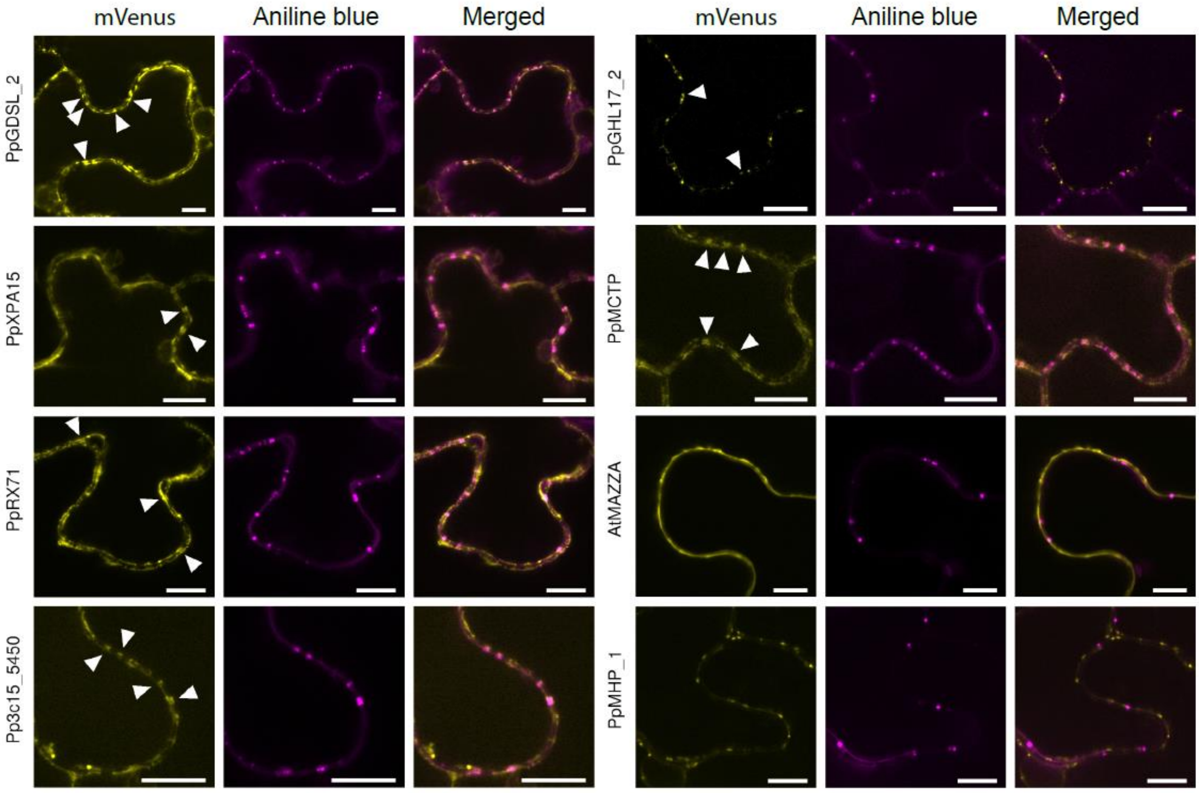


**Supplementary Figure S5:** Subcellular localization of a selection of representative candidates from *P. patens* PD proteome in *N. benthamiana* leaf epidermis cells. Single optical sections at cell-to-cell interface showing the co-localization of selected candidates with plasmodesmata marker aniline blue. AtMAZZA and PpMHP_1 did not overlay with PD marker and were scored with the PD index as non-PD localized proteins (Figure 3B). Scale bars, 10 µm.

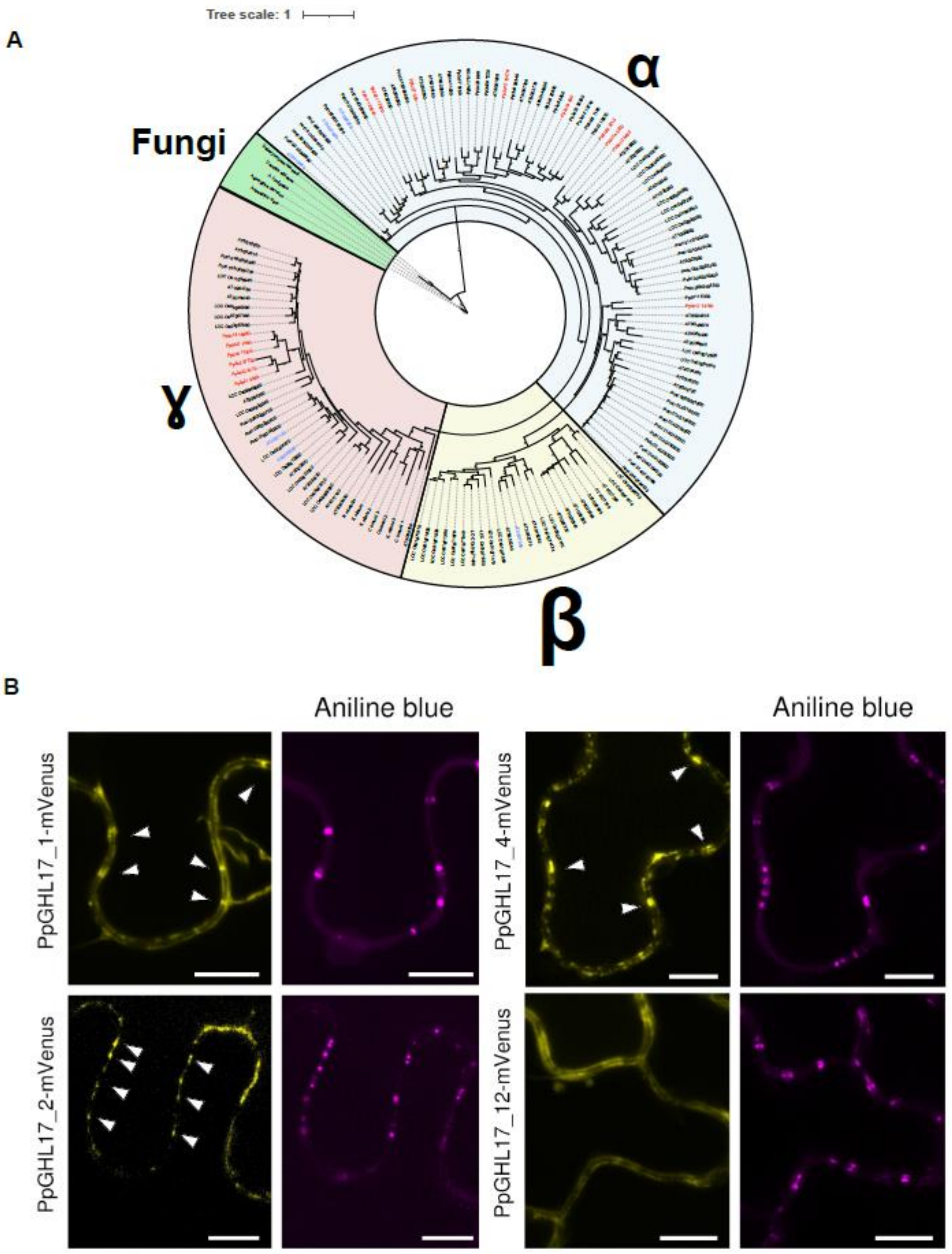


**Supplementary Figure S6:** PD localization of GHL17 proteins correlates with phylogenetic distribution. (**A**) Phylogenetic tree including all GHL17 sequences used in the phylogenetic analysis (Supplementary Table 5). *P. patens* GHL17 proteins identified in the PD proteome are labeled in red and Arabidopsis orthologs previously assayed for localization colored in blue (1). $\alpha$ and β clades are shadowed in yellow and blue, respectively. Support values from 1000 bootstrap samples are shown at the nodes. (**B**) Subcellular localization of *P. patens* GHL17s from alpha (PpGHL17_1, PpGHL17_2 and PpGHL17_4) and gamma (PpGHL17_12) clades in *N. benthamiana* leaf epidermis cells. Aniline blue was used as PD marker. Scale bars, 10 µm.

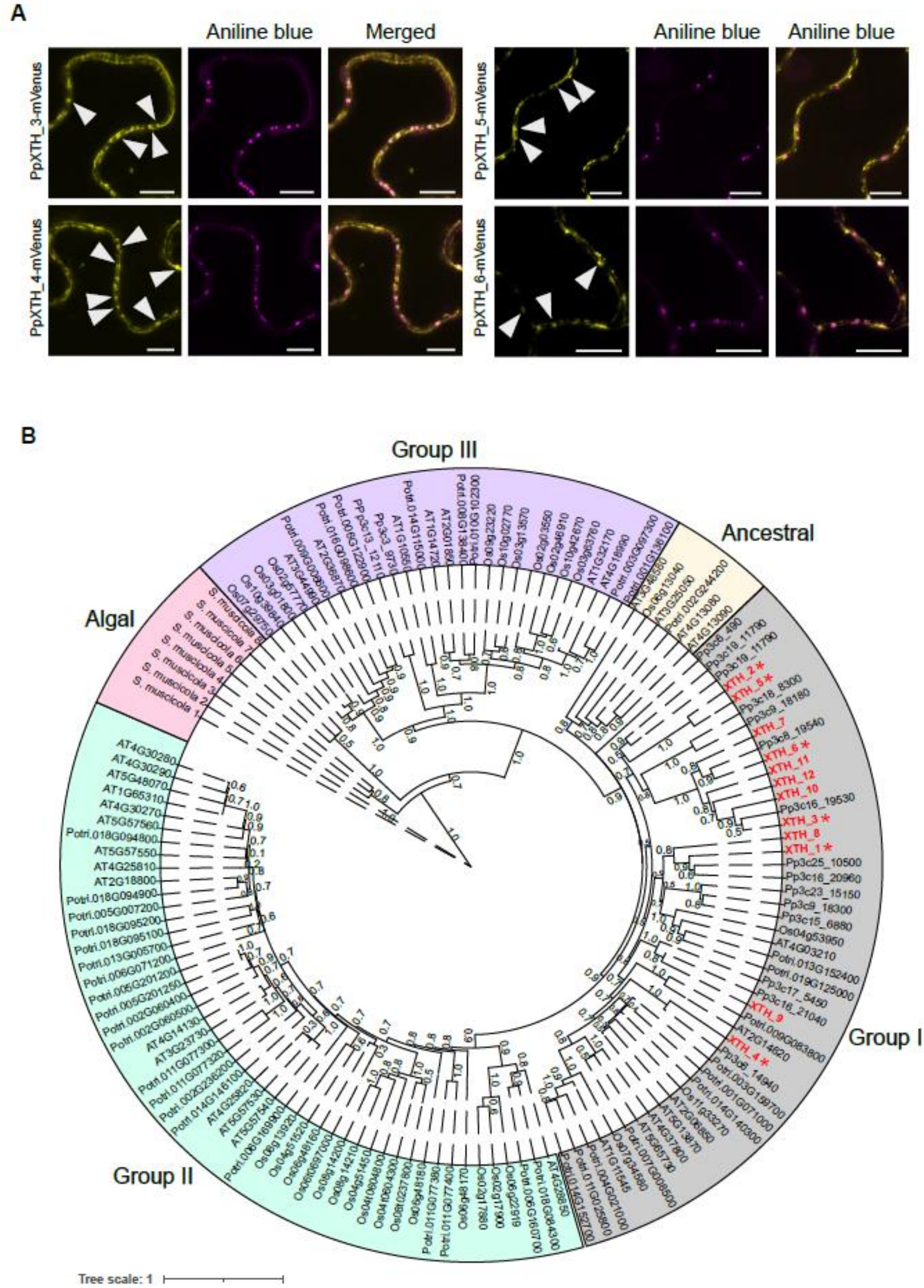


**Supplementary Figure S7**: PD-localized *P. patens* XTHs grouped in cluster I. (**A**) Subcellular localization of PpXTHs in *N. benthamiana* leaf epidermis cells showing overlay with aniline blue. Scale bars, 10 µm. (**B**) Phylogenetic tree of Arabidopsis, *O. sativa*, *P. trichocarpa*, *P. patens* and *S. muscicola* XTH proteins. *P. patens* XTH proteins identified in the HC300 PD proteome are labeled in red and clustered in group I. Red asterisks indicate *P. patens* PD-localized XTH proteins. Support values from 1000 bootstrap samples are shown at the nodes.

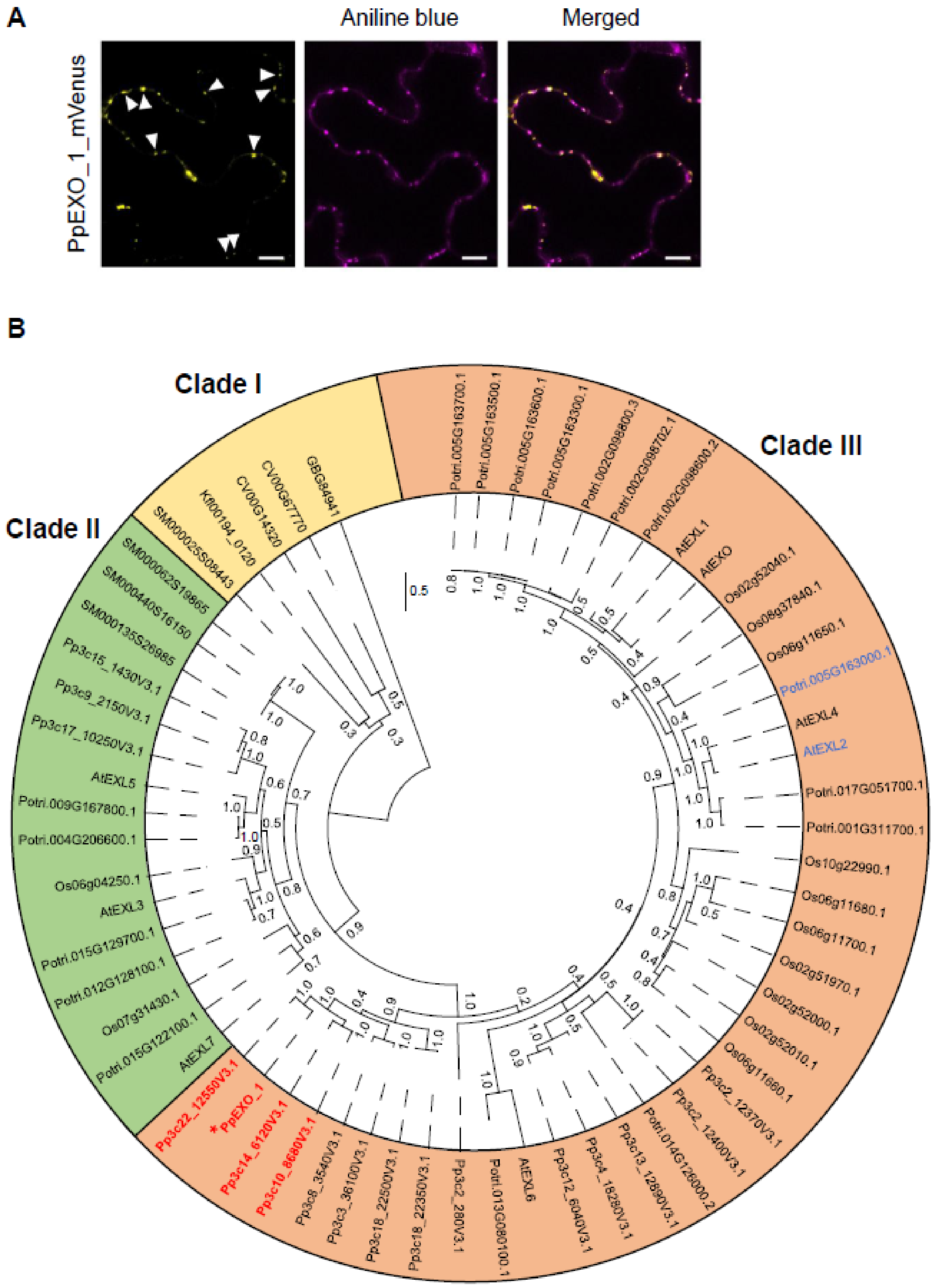


**Supplementary Figure S8**: EXO/EXL family members. (**A**) Subcellular localization of PpEXO_1 (Pp3c19_8770V3.5.p) in *N. benthamiana* leaf epidermis cells showing overlay with aniline blue. Scale bars, 10 µm. (**B**) Phylogenetic tree of EXO/EXL family members from representative species of the embryophytes (17 from *P. patens*, 8 from Arabidopsis, 12 from *O. sativa,* and 17 from *P. trichocarpa*) and all algae for which EXO orthologs could be identified (*Coccomyxa subellipsoidea*, *Klebsormidium nitens*, *Chara braunii*, *Spirogloea muscicola*). *P. patens* EXO/EXL members identified in the HC300 PD proteome are labeled in red and clustered in clade III in which EXO identified in other PD proteomes are grouped (labeled in blue).

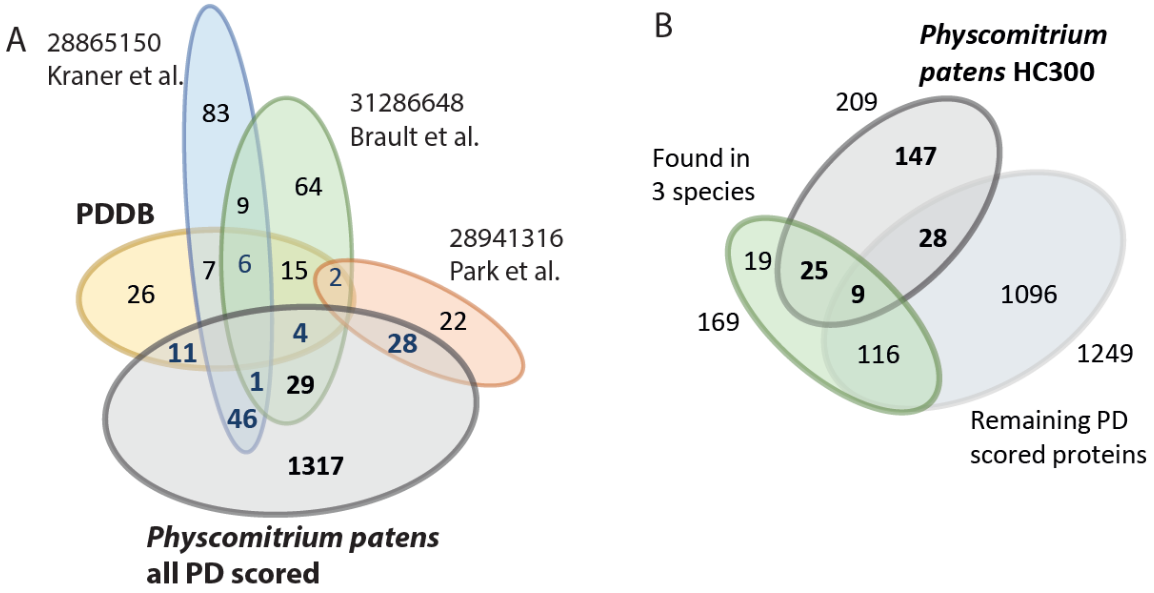


**Supplementary Figure S9**: Overlap of *Physcomitrium* PD proteomes with the *bona fide* set of 338 proteins from different literature sources. (**A**) Venn diagram including all PD scored proteins. (**B**) Venn diagram including HC300, the remaining PD scored proteins, and proteins identified in PD proteomes of at least 3 species (see Figure 1F).

### SI Materials and Methods

*Growth conditions for Physcomitrium – Physcomitrium patens* ecotype Gransden cultures were obtained from the *International Moss Stock Center (IMSC)* (Freiburg). *P. patens* was cultured on modified BCD medium containing 5 mM diammonium tartrate (2) in Petri dishes at 21°C under long day conditions (16 h light/8 h dark) at ∼90 µmol $m^{-2}$ $sec^{-1}$, 60% humidity. To avoid precipitation of the modified BCD medium, the components were prepared in separate solutions which were mixed directly before use: Solution B (0.1 mM $MgSO_4 \cdot 7H_2O$), Solution C (1.84 mM $KH_2PO_4$), Solution DK (1 M $KNO_3$), Solution DF (4.5 mM $FeSO_4 \cdot 7H_2O$), Trace Element Solution (TES) (614 mg $H_3BO_3$, 55 mg $CuSO_4 \cdot 5H_20$, 28 mg KBr, 110 mg $Al_2(SO_4)3 \cdot K_2SO_4 \cdot 24H_20$, 389 mg $MnCl_2 \cdot 4H_20$, 55 mg $ZnSO_4 \cdot 7H_20$,LiCl, 10 mM $CoCl_2 \cdot 6H_20$, 28 mg KI and 28 mg $SnCl_2 \cdot 2H_20$ in 1 L $H_20$), 500 mM diammonium tartrate and 1 M $CaCl_2$ Solution were prepared separately, autoclaved and stored under sterile conditions. To prepare the modified BCD directly for use, 10 mL each of solutions B, C, DK, DF and TES, and 8 g agar were mixed, made up to a volume of 1 L with sterile water, adjusted to a final pH of 5.4 and autoclaved. Finally, 1 mL 1 M $CaCl_2$ was added to complete the BCD medium.

For liquid cultures, the same medium was used without the addition of agar, pH was left at 6.5 and 1% (m/v) Glucose was added before autoclaving. Either moss from plates or moss from liquid cultures was homogenized using an ULTRA-TURRAX and then transferred to 100 ml medium in a 250 ml flask. Moss was cultured for two weeks before harvest under the same conditions as described above, with one medium exchange after one week. Harvested moss samples were kept at -80°C.

*Growth conditions for Nicotiana benthamiana – N. benthamiana* seeds were soaked in water for 24 h at 20°C and then placed in peat moss substrate for germination in a growth chamber. After 2 weeks, uniform seedlings were chosen and transferred to a greenhouse for 3 weeks under long day conditions. Day and night temperatures in the greenhouse were maintained at 22 and 20°C, under long day conditions (16 h light/8 h dark) at ∼90 µmol $m^{-2}$ $sec^{-1}$, 60% humidity.

*Plasmid construction* – Transient expression *N. benthamiana*: For the cloning of 149 *P. patens* open reading frames (ORFs) into the binary pAB134-mVenus plasmid (3), total RNA was extracted from *P. patens* protonemal and gametophore tissues using RNeasy plant mini kit (Qiagen), followed by DNase I treatment according to the manufacturer instructions. The poly(A)+ fraction was purified using DynaBeads Oligo(dT)25 (Thermofisher) and cDNA was synthesized using the Maxima H Minus First Strand cDNA synthesis kit (Thermofisher). ORFs from PD candidates were amplified in a 2-step PCR reaction to attach the GATEWAY attB sites (primers listed in Supplementary Table 6). Purified PCR products were cloned into pDONR221 with a BP clonase reaction (Thermofisher). *E. coli* Stellar cells (Takara Bio) were transformed with the resulting entry vectors, and vector identity and integrity were confirmed after DNA sequence analysis using M13 forward and M13 reverse primers for complete coverage of the insert sequence. Plasmids with the correct sequence were isolated and recombined by LR reaction into the binary plant expression vector pAB134-mVenus (3) as N-terminal mVenus fusions under control of a β-estradiol-inducible promoter. Stable expression in *P. patens*: To generate pTH-PpGHL17_1-mVenus construct, the genomic sequence of PpGHL17_1 (Pp3c17_13760) was amplified from genomic DNA isolated from 7-day-old protonemal tissue with primers containing the regions of homology to the attL1 site of pDONR221 vector and the 5´end of mVenus ORF. pDONR221 was amplified with primers harboring the attL1 and attL2 sites and mVenus ORF was amplified with primers containing the 3´ end and attL2 site of PpGHL17_1 and pDONR221, respectively. The PCR products were gel purified, mixed 1:1:1 molar ratio and incubated with In-fusion reaction mix (Takara). Subsequently, PpGHL17_1-mVenus was subcloned into the pTH Ubi-Gate (4) expression vector by LR reaction.

*Transient gene expression in Nicotiana benthamiana* – We employed *Agrobacterium tumefaciens* strain GV3101 pMP90 containing an expression cassette for the p19 silencing suppressor from the tomato bushy stunt virus to enhance efficiency of transgene expression (5). Agrobacteria were transformed with binary plant expression plasmids by heat-shock and plated onto Luria-Bertani (LB) agar plates (10 g/L tryptone, 5 g/L yeast extract, 10 g/L NaCl, bacto agar 15 g/L) supplemented with appropriate antibiotics (50 µg/mL rifampicin, 50 µg/mL gentamicin, 50 µg/mL kanamycin and 100 µg/mL spectinomycin). Single Agrobacterium colonies were used to inoculate 5 mL dYT medium (16 g/L tryptone, 10 g/L yeast extract, 5 g/L NaCl) or LB broth (10 g/L tryptone, 5 g/L yeast extract, 10 g/L NaCl) supplemented with antibiotics as above and were incubated at 28°C overnight in a shaking incubator. To ensure that Agrobacteria were in the exponential growth phase, 0.5 mL of overnight culture was used to inoculate 5 mL fresh medium supplemented with antibiotics, and bacteria were cultured for 5-6 hours to reach $OD_{600}$ ≅0.6. Bacteria were then sedimented and resuspended in Infiltration Buffer (5% sucrose, 10 mM $MgCl_2$, 450µM acetosyringone, 10mM MES pH 5.6) and adjusted to $OD_{600}$ 0.3. The PM-localized receptor-like kinase MAZZA fused to mCherry (6) was co-infiltrated at $OD_{600}$ 0.1. Leaves from 3-week-old *N. benthamiana* plants were infiltrated on the abaxial leaf surface using a 1ml syringe without a needle. Two days after infiltration, gene expression was induced by spraying infiltrated leaves with β-estradiol solution (10 µM β-estradiol, 0.1% v/v Tween-20, (7)) and plants were grown under continuous light at 21 °C. Induction time was dependent on protein maturation times and ranged from 2 to 24 h.

*Transformation of Physcomitrium via protoplast transformation* – Protoplasts were transformed using a PEG-mediated protocol as described (8). Briefly, protoplasts were isolated from 6-7 day old protonemal tissue using 2% Driselase (Sigma) in 8.5% (w/v) D-mannitol. pTH-GHL-mVenus plasmid was linearized with *Swa*I and 15µg plasmid was used for transformation of protoplasts (1.2 x $10^6$ protoplasts/ml). Protoplasts were resuspended in Protoplast Regeneration Medium Top Layer (PRMT: BCD medium, 8 (w/v) D-mannitol, 5mM diammonium tartrate, 10mM $CaCl_2$, 0.7% (w/v) agar), then plated and regenerated on Protoplast Regeneration Medium Bottom Layer (PRMB: BCD medium, 6 (w/v) D-mannitol, 5mM diammonium tartrate, 10mM $CaCl_2$, 1% (w/v) agar) on top of a cellophane sheet to enable transfer between selective and non-selective media. After 4 days on PRMB, the cellophane sheet was transferred to BCD + $NH_4$-tartrate plates supplied with 10 µg/ml hygromycin for selection.

*Total protein extraction – Physcomitrium patens* tissue was finely ground in liquid nitrogen. 50-200 mg of power was mixed with 0.5-1 ml extraction buffer (10 mM Tris-HCl pH 8.0, 6 M urea, 2 M thiourea, 5 mM DTT, 1 mM PMSF, 2% w/v PVPP, 0.5% v/v Protease inhibitor cocktail (P9599, Sigma-Aldrich) (9). The sample was then incubating vigorously for 1h at 4 °C, and then centrifuged at 12,000 x g for 10 min at 4 °C. The total proteins solubilized in the supernatant was transferred to a fresh tube and precipitated in 80% acetone at -20 °C overnight. Afterwards, the proteins were pelleted at 12,000 x g for 10 min at 4 °C and washed with 1 ml ice-cold 80% acetone twice. The pellet then was solubilized in a 100-300 µl resuspension buffer (10 mM Tris-HCl pH 8.0, 6 M urea, 2 M thiourea).

*Microsomal fraction preparation* – The microsomal fraction isolation was carried out according to (10) with small modifications. A total of 1 to 1 g of *Physcomitrium patens* tissue was homogenized with 10 ml ice-cold extraction buffer (50 mM Tris-MES pH 7.5, 330 mM mannitol, 100 mM KCl, 1 mM EDTA, 5 mM DTT, 1 mM PMSF, 0.5% v/v Protease inhibitor cocktail (P9599, Sigma-Aldrich). The homogenate was centrifuged for 20 minutes at 8000 × g at 4 °C, and the supernatant was centrifuged for 75 minutes, 48,000 × g at 4 °C to make the microsomal fraction pellet. The microsomal fraction pellet was resuspended in 100 µl of 25 mM Tris-MES pH 7.5, 330 mM Sucrose, 0.5 mM DTT.

*Cell wall fraction and plasmodesmata fraction preparation* – One gram of starting material was homogenized in a glass potter grinder with 10ml ice-cold extraction buffer (100 mM Tris-HCl pH 8.0, 150 mM NaCl, 10 mM EDTA, 1 mM DTT, 1% Triton X-100, 0.5% sodium deoxycholate, 0.1% SDS, 10% glycerol, 0.5% v/v protease inhibitor (P9599, Sigma-Aldrich). The homogenate was centrifuged for 10 minutes at 1000 × g at 4 °C and washed (washing buffer: 100 mM Tris-HCl pH 8.0, 150 mM NaCl, 10% glycerol, 0.1% Triton X-100) four times. The resulting pellet containing cell wall and tissue debris was then ground into fine powder in liquid nitrogen to further eliminated organelle contaminations and prepare the cell wall for digestion with Driselase, by breaking up remaining cell clusters (11). The ground sample was fully mixed and incubated for 20 min on ice, then it was centrifuged with a swing-out rotor for 15 min, 1500 x g at 4 °C and the supernatant was discarded. The grinding step was repeated one time. Then the pellet was washed with 10 ml wash buffer (100 mM Tris-HCl pH 8.0, 150 mM NaCl, 10% glycerol, 0.1% Triton X-100) and centrifuged for 10 min, at 1500 x g at 4 °C. To enrich the plasmodesmata fraction, the cell wall pellet was subjected into 10 ml cell wall digestion buffer (10 mM MES-KOH, pH 5.5, 4.4% mannitol, 0.7% w/v of Driselase (D8037, Sigma-Aldrich) and digested for 1.5 h at 37 °C, 100 rpm shaking. To remove the undigested cell wall fragment, the sample was centrifuged with a swing-out rotor for 15 min, 1500 x g at 4 °C. The supernatant was taken to ultracentrifugation (No. 344059, Beckman), 110,000 x g for 1 h at 4°C. The pellet was washed with ice-cold TBS buffer (20 mM Tris-HCl pH 7.4, 140 mM NaCl, 2.5 mM KCl) and resuspended in a minimal volume of resuspension buffer (10 mM Tris/HCl pH 8, 6 M Urea, 2 M Thiourea).

*Protein clean-up, trypsin digestion and peptide desalting* – The single-pot, solid-phase-enhanced sample-preparation (SP3) technology (12) follow-up with trypsin digestion was applied to get rid of the detergent in samples for proteomics. Reduction of disulfide bridges was obtained by an excess of DTT (6.5mM) and alkylation of free cysteines was done by iodoacetamide (27 mM). SpeedBead Magnetic Carboxylate Modified Particles (SP3 beads, 45152105050250 / 65152105050250, GE Healthcare) were pre-washed by Milli-Q water. After adjusting concentration of the SP3 beads to 50µg/µl in Milli-Q water, they were added into the protein sample according to a ratio of 1:10 protein:beads. 50% Ethanol was then introduced in the sample for binding induction. After incubating 15 min at 24°C, 1,000 rpm shaking, the beads were washed three times by 80% Ethanal and subjected to trypsin digestion buffer (100 mM Ammonium bicarbonate, trypsin (trypsin: protein ratio 1:100, V5113, Promega) for 12-14 h at 37°C. Digested proteins were acidified to pH 2 using TFA and were then desalted over C18 STAGE tips (13).

*LC-MS/MS analysis of peptides* – Peptide mixtures were analyzed by nanoflow Easy-nLC (Thermo Scientific) and Orbitrap hybrid mass spectrometer (Q-exactive HF, Thermo Scientific). Peptides were eluted from a 75 µm x 25 cm analytical $C_{18}$ column (PepMan, Thermo Scientific) on a linear gradient running from 5% to 90% acetonitrile for 70 min. Proteins were identified based on the information-dependent acquisition of fragmentation spectra of multiple charged peptides. Up to ten data-dependent MS/MS spectra were acquired for each full-scan spectrum acquired at 60,000 full-width half-maximum (FWHM) resolution.

*Peptide and protein identification* – Protein identification and ion intensity quantitation was carried out by MaxQuant version 2.0.3.0 (14). Spectra were matched against the *Physcomitrium patens* proteome (Ppatens_318_v3.3.protein.fasta, 87533 entries) using Andromeda (15). Thereby, carbamidomethylation of cysteine was set as a fixed modification; oxidation of methionine as well as phosphorylation of serine, threonine and tyrosine were set as variable modifications. Mass tolerance for the database search was set to 20 ppm on full scans and 0.5 Da for fragment ions. Multiplicity was set to 1. For label-free quantitation, retention time matching between runs was chosen within a time window of two minutes. Peptide false discovery rate (FDR) and protein FDR were set to 0.01, while site FDR was set to 0.05. Hits to contaminants (e.g. keratins) and reverse hits identified by MaxQuant were excluded from further analysis. The mass spectrometry proteomics data have been deposited to the ProteomeXchange Consortium via the PRIDE partner repository (16) with the dataset identifier PXD032820.

*Confocal fluorescence microscopy* – Live confocal images were obtained using either a Zeiss LSM 900 or an Olympus SpinSR10 microscope. The Zeiss LSM 900 was equipped with Airyscan GaAsP-PMT detectors and diode lasers using a C-Apochromat 40x/1.20 W Korr FCS water objective. The pinhole size was set to 1 AU. To visualize proteins of interest, mVenus was excited with a 488 nm laser at 1-4% laser power and fluorescence emission was detected at 520-579 nm. To visualize the MAZZA-mCherry PM marker (6), mCherry was excited at 561 nm with 2-5% laser power and emission was detected at 580-617 nm. To avoid chloroplast autofluorescence, both lasers were blocked by a 620 nm short-pass filter. To visualize pit fields, *N. benthamiana* leaves were infiltrated with 0.1% (w/v) aniline blue and imaged immediately (excitation at 405 nm and 3-5% laser power, emission detected at 440-480 nm). For colocalization, mVenus, mCherry and aniline blue channels were set as separate tracks to minimize cross-talk and images were acquired using sequential line scanning. The Olympus SpinSR10 was equipped with a 40x objective with SIL300CS-300CC Silicone immersion oil, CCD camera and excitation lasers at 405 nm for aniline blue and 514 nm for mVenus. Confocal images were deposited in the Bioimage Archive (https://www.ebi.ac.uk/bioimage-archive/) under accession number S-BIAD466.

For moss, protonema and gametophore cells expressing GHL17-mVenus (Pp3c17_13760) were incubated in 0.01% (w/v) aniline blue for 20 min and washed twice with a modified BCD liquid medium prior to imaging. Samples were sequentially excited at 405 nm, with 7-10% laser power and emission detected at 440-480 nm for aniline blue, and at 488 nm, 0.5-2% laser power, and emission wavelength of 520-580 nm for mVenus. Raw Airyscan images were processed in Zen software (Zeiss) using deconvolution mode at default settings.

*Quantification of PD candidate localization in N. benthamiana* – Confocal images were initially screened visually for colocalization of the protein of interest and aniline blue. Candidates were first scored based on the number of times the YFP signal overlaid with aniline blue (*i.e.*, colocalized) per cell (Supplementary Table 3) to differentiate between candidates localizing to PD and those not. Candidates showing colocalization with aniline blue were selected for further analysis to determine the degree of enrichment at of the protein of interest at the PD compared to other regions of the plasma membrane, the so-called PD index (17). To minimize bias in this quantification, we used a semi-automated approach based on an in-house-generated macro for the *Fiji* software package (18). In brief, PD index values were calculated by dividing the candidate mVenus-signal intensities found in regions of interest (ROIs) of PDs, divided by the mean signal intensities measured in background ROIs at the plasma membrane, not associated with PDs. Thus, PD index values higher than the negative controls indicate accumulation of the candidate protein at PD sites compared to the rest of the plasma membrane. PD-ROIs were centrally placed on the thresholded strong aniline blue signal and background ROIs were placed on the centered line at the plasma membrane after processing the MAZZA-mCherry or the weaker background aniline blue signal. Both types of ROIs were generated with the same size based on the average size of thresholded PD found in the particular image. Code of the Fiji macro can be accessed at: https://github.com/SHaensch/2022_PD-index-quantification. GraphPad Prism software was used to generate the box plots.

*PD index quantification* – For PD index quantification, fluorescence microscopy measurements were processed using an in-house written *Fiji* macro (18). The macro allows to crop regions with the most clear PD sites. Subsequently channels of aniline blue, the candidate YFP signal and, if applicable, PM marker MAZZA-mCherry, were separated. PDs were identified by an auto-threshold on the aniline blue channel according to the *Fiji YEN*-algorithm (19) and modified by the user to the best fit, if necessary.

To ensure reproducibility, threshold-values were saved and automatically documented along with PD index results later on. Center coordinates of PD were identified and at each PD position a square ROI was set at the length size of the square root of the average PD area. To define background ROIs, the weak signal of the aniline blue channel or MAZZA-mCherry as PM marker were used. Previously identified regions of PDs were automatically excluded from background ROI placement. A combination of *median filtering, local thresholding*, smoothing by *dilation* and *erosion* of the binary masks, *skeletonization* of the resulting masks and finally applying the *ultimate points* function (returning the coordinates of the ultimate eroded points of the euclidean distance map of the binary thresholded images) was used for placing background ROIs. These steps positioned the background ROIs in the centered line of the cell membrane regions with the same size as PD-ROIs of the previous steps. To automatically fit the size of PD and background regions best, parameters were modified when using measurements of the Olympus spinning disc microscope (e.g. PD ROI and background ROI size was enlarged by 50% due to different sampling rate and radius of median filtering, local thresholding and smoothing factors due to different signal intensities). Per image, the PD index was calculated as the original candidate YFP mean intensity values of all PD ROIs divided by those of background ROI values. Thus, PD index values higher than the negative controls indicate accumulation of the candidate protein at PD sites compared to the rest of the plasma membrane. Along with the numerical results (PD-index, thresholds and number of PDs), visual outputs were generated to allow for subsequent visual quality control of the quantification e.g. placement of all ROIs and generated masks.

*Transmission electron microscopy* and PD quantification – Protonema tissue grown as liquid cultures were processed following the protocol described in (20) with slight modifications. Briefly, the tissues were fixed in 2.5% (v/v) glutaraldehyde (Agar Scientific, R1020) in 0.1 M sodium cacodylate (Agar Scientific, R1103) buffer, pH 6.9, for 2h at room temperature, then at 4°C overnight. Subsequently, samples were rinsed in 0.1M sodium cacodylate buffer and post-fixed with 2% (w/v) osmium tetroxide (EMS, E19150) supplemented with 0.15% potassium ferricyanide (Sigma-Aldrich, 60299) in water for 2 hours at 4°C. After thorough rinsing first with buffer then with water, the material was gradually dehydrated through an ethanol series and transferred to acetone. Infiltration with and embedding into Araldite 502/Embed 812 resin (EMS, 13940) was done with the help of an EMS poly III (EMS, 4444). Ultrathin sections (70-90 nm) were produced using a PowerTome PTPC (RMC Boeckeler) and collected on nickel slot grids as described by Moran and Rowley (21). After staining with 0.1 % potassium permanganate (Merck KGaA, 105082) in 0.1 N $H_2SO_4$ (22), either alone or followed by 2% aqueous uranyl acetate (TAAB, U007), sections were examined with a Hitachi H-7650 transmission electron microscope (Hitachi High Technologies Europe) operating at 100 kV.

PD density was calculated by counting PD number per µm of cell-to-cell interface length as measured by Fiji and converted to number per $\mu m^2$ of cell wall using the Gunning constant, 1/(T + 1.5R), where T is section thickness (nm) and R is the average radius of PD (nm) (23, 24). The average thickness of the ultrathin section was 80 and average radius of PD was 12.5 nm. To avoid double counting, no directly consecutive sections were analyzed.

*Cryo transmission electron microscopy* – Pellets of the plasmodesmata fraction preparation were resuspended in 150 mM PBS pH 7.95, 0.1% (v/v) DDM. R2/1 Cu200 mesh copper EM grids (Quantifoil R2/1 Cu 200 holey carbon) were glow discharged for 30 s and 4 µl of resuspended sample were deposited on the grid and plunge frozen in liquid ethane/propane using a Vitrobot Mark 4 system (FEI) at 100% relative humidity and 6°C. Cryo grids were then stored in liquid nitrogen for further processing. Images were acquired on a Titan Krios 300 kV (FEI) transmission electron microscope (TEM) equipped with a post column energy filter (Gatan) operated in zero-energy-loss mode with a slit width of 20 eV, and K2 direct electron detector (Gatan) in counting mode. SerialEM (bio3d.colorado.edu/SerialEM/) and Digital Micrograph (Gatan) software were used for data acquisition. Dose-fractionated movie frames were recorded under a 100 µm objective aperture at a magnification of 42,000x (3.52 Å per pixel) with an exposure time of 8s amounting to a total exposure of 60 e-/Å2 and target defocus -5 µm. Motions between individual frames were corrected by MotionCor2 package.

*Phylogenetic analysis* – Protein sequences of GHL17s, XTHs and EXOs were retrieve from public databases (8, 20, 21, 25, 26) and were compiled in Supplementary Table 5. All isolated GHL17 amino acid sequences were subjected to protein domain prediction at InterPro server (27) and only those sequences containing the glycoside hydrolase, family 17 domain were used in the phylogenetic analysis (Supplementary Table 5). Amino acid sequences from 151 GHL17 orthologs were aligned using the MAFFT method with a gap extension penalty of 0.123 and 1.53 of gap opening penalty and curated with BMGE. Phylogenetic tree was generated using the NGPhylogeny.fr tool (28) and visualized and edited in iTOL (29). The lower BIC (Bayesian Information Criterion) model was selected in PhyML (30) for each tree. Therefore, LG+G+I model was used for the GHL17s and XTHs trees while the WAG+G model was used for the EXOs tree. Statistical branch support was calculated using 1000 bootstrap replications.

*Structural analysis* – Using a full local installation of AlphaFold 2.2.0, structural models for all 16 *Pp*GHL17s found in the proteome were generated (31). Models were selected for highest average confidence scores (pLDDT) and corresponding structure predictions were aligned and rendered in ChimeraX 1.4 (32). Potential GPI anchor attachments at C-terminal regions were predicted using PredGPI (33).

*CFDA transport assay* – To assess transport through PD, CFDA solution in BCD medium was prepared immediately before use. Protonema cells were incubated with the 50 µM CFDA and kept under light (21 °C) for 30 minutes. After introducing CFDA to plants, acetated groups of CFDA are cleaved by steranes to convert CFDA to CF which is no longer plasma membrane-permeable and it moves through PD. After rinsing the stained protonema tissues three times with distilled water, protonema cells were observed under a Zeiss LSM 900 microscope with a C-Apochromat 40x/1.20 W Korr FCS water objective 40X, excitation at 488 nm, and 525/50 nm emission. The fluorescent signal was bleached in a single cell with a 405 nm, with 60% laser power and its recovery was monitored within 15 minutes. The detected signals of 6 independent experiments were densitometrically quantified using Fiji-3 imaging software.

*Statistical analyses and data visualization* – Functional classification of proteins was done based on MAPMAN (34). Information about subcellular location was derived from SUBA (35). Detailed protein function was manually updated with the support of TAIR (36). Other statistical analyses were carried out with Sigma Plot (version 11.0) and Excel (Microsoft, 2013). Over-representation analysis was done via Fisher's exact test, p values were adjusted using Bonferroni correction. Interactome data from published large-scale data sets were obtained from Arabidopsis Interactome Viewer 2 (37) or STRING (38).
