## Supplementary figures and images for "A high confidence *Physcomitrium patens* plasmodesmata proteome by iterative scoring and validation reveals diversification of cell wall proteins during evolution"

### Supplementary Figure S1_PD prep workflow

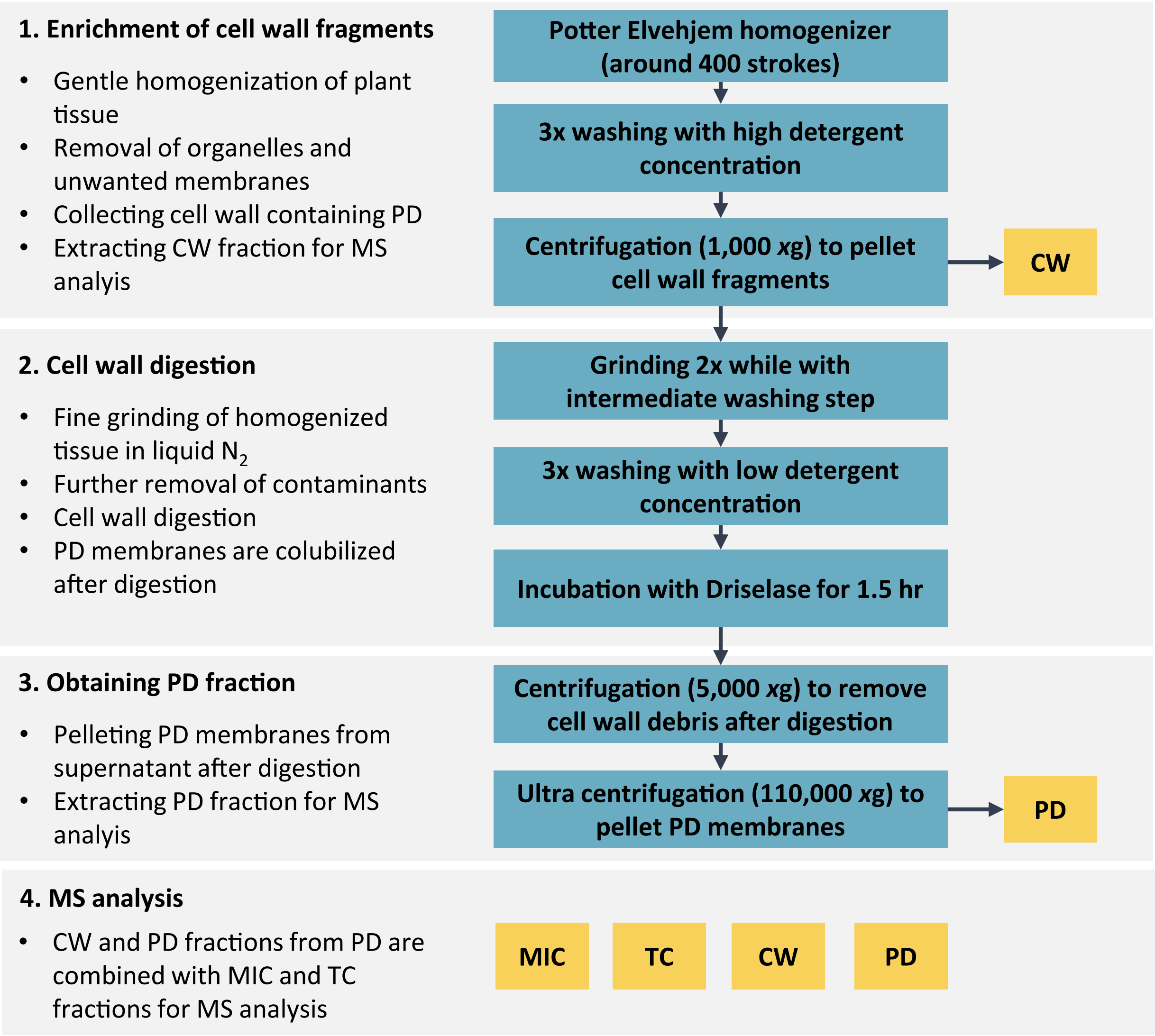


Supplementary Figure S1

### Supplementary Figure S2_TEM

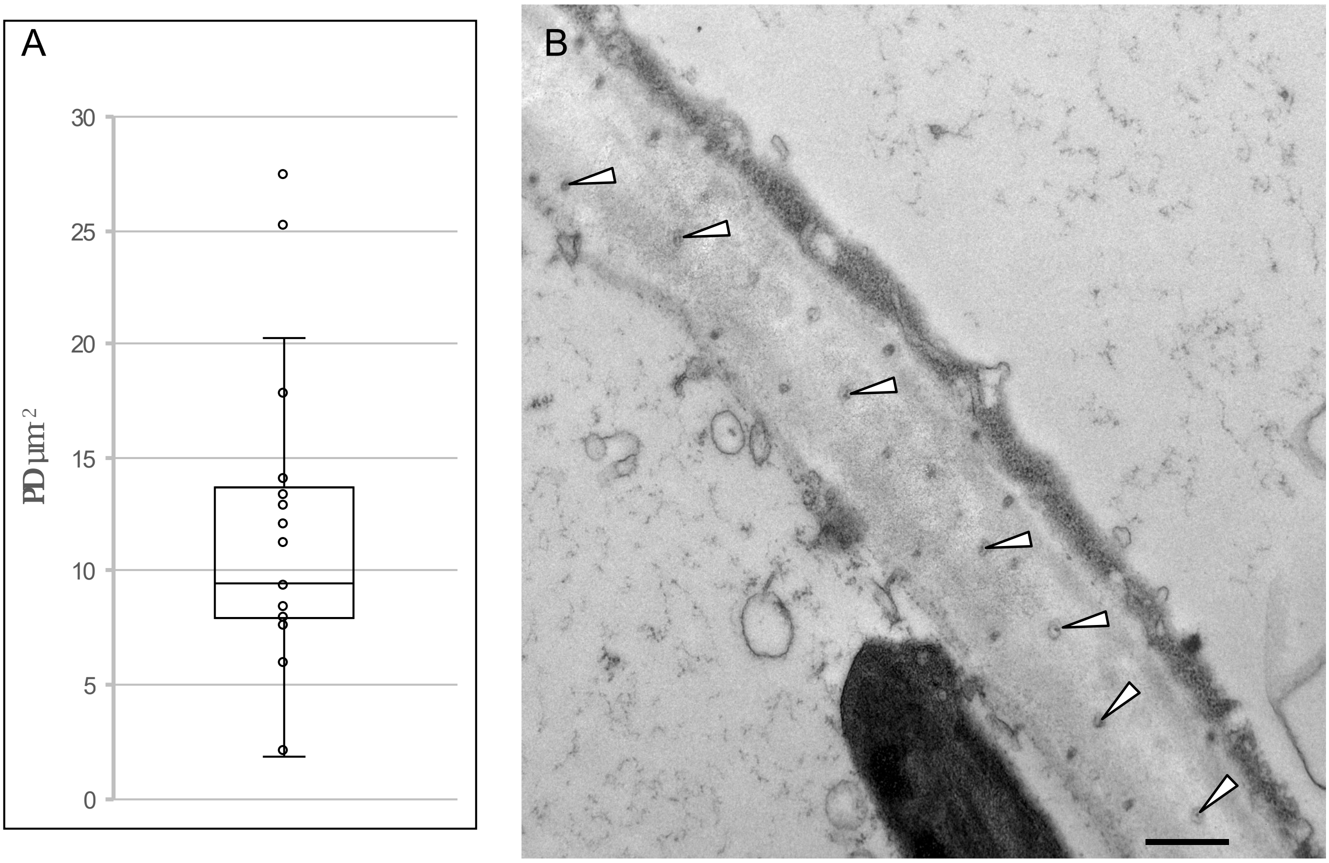


Supplementary Figure S2

### Supplementary Figure S3_PDDB

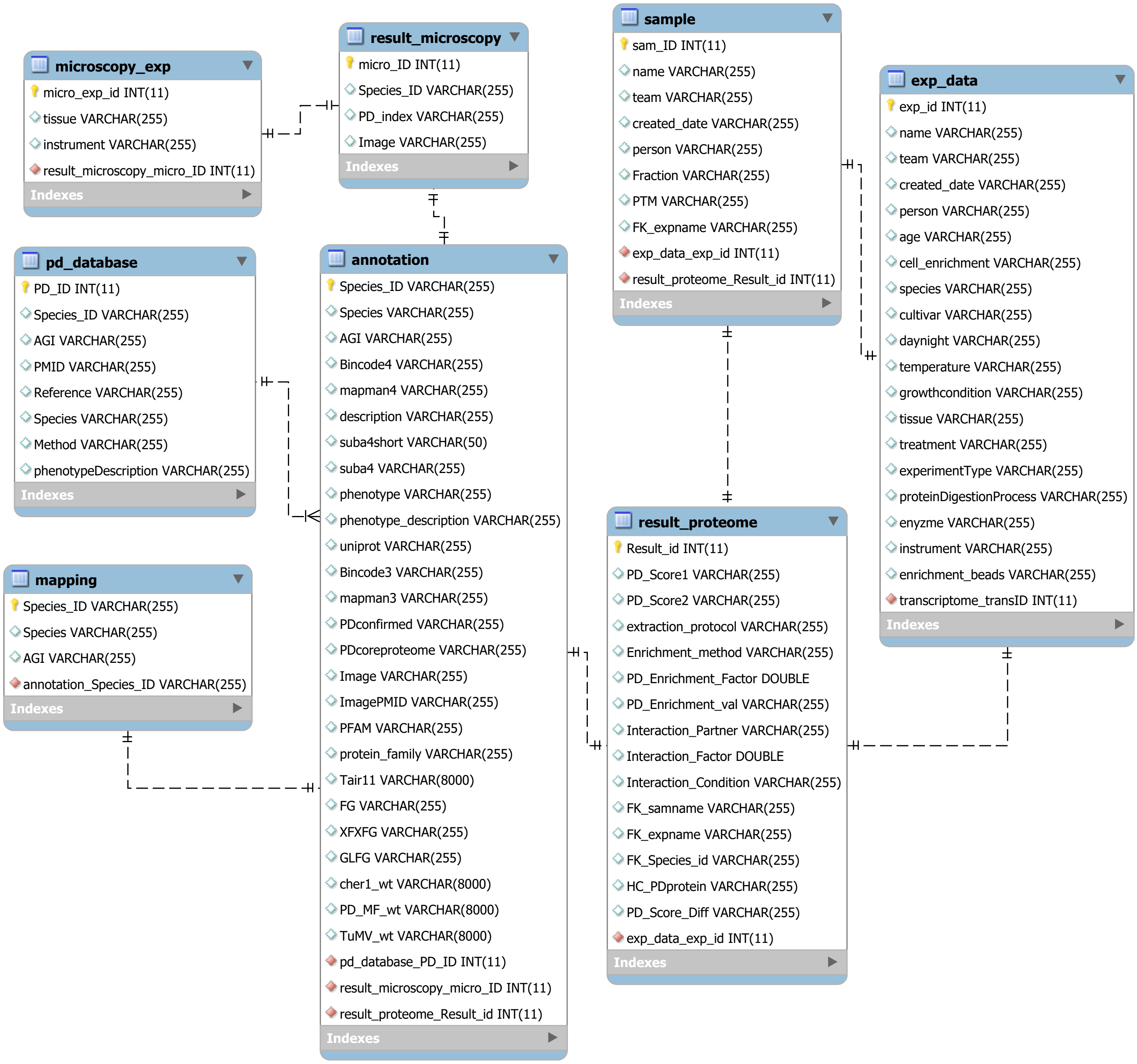


Supplementary Figure S3

### Supplementary Figure S4_PD scoring formeln

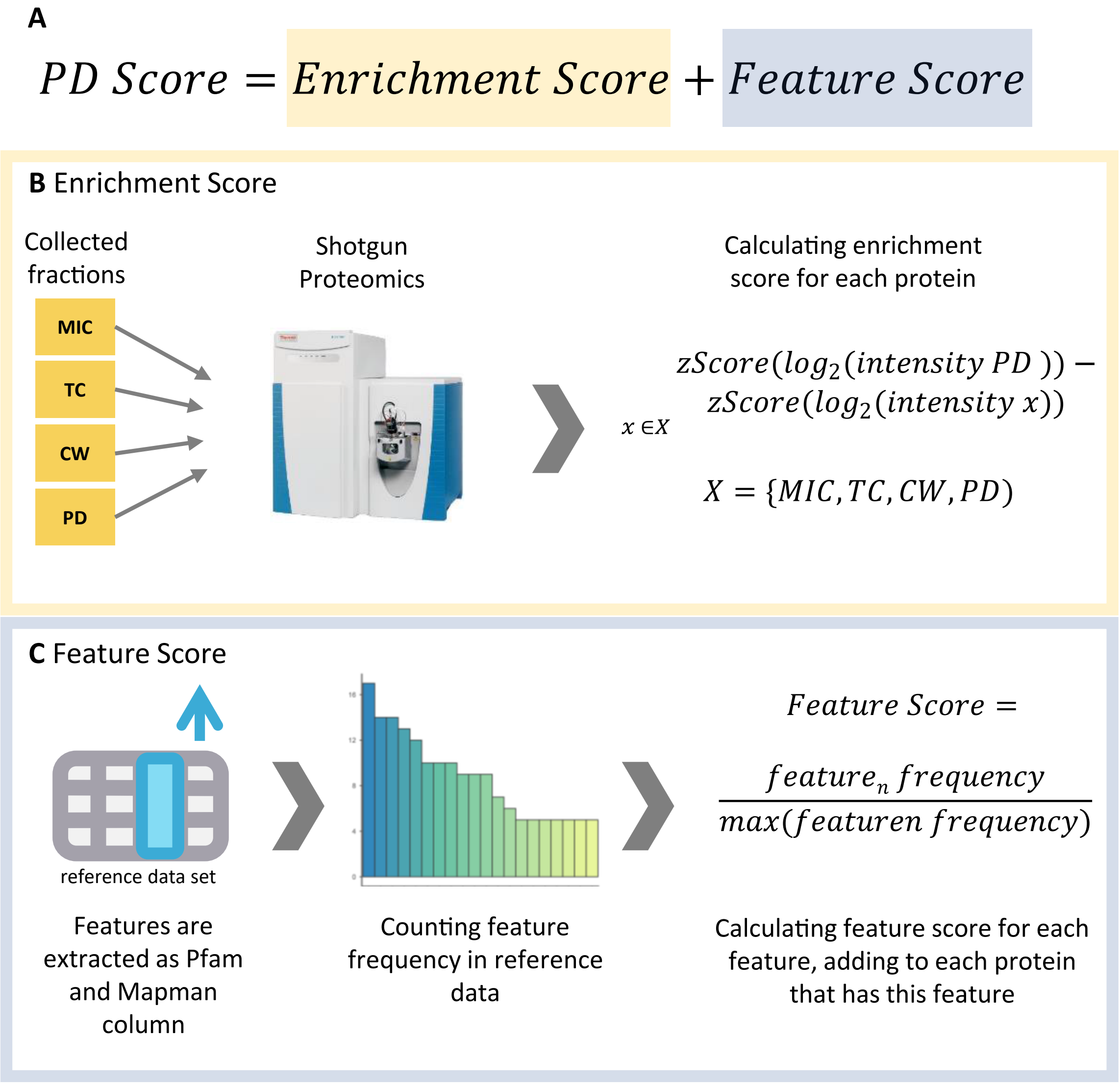


Supplementary Figure S4

### Supplementary Figure S5_more localizations

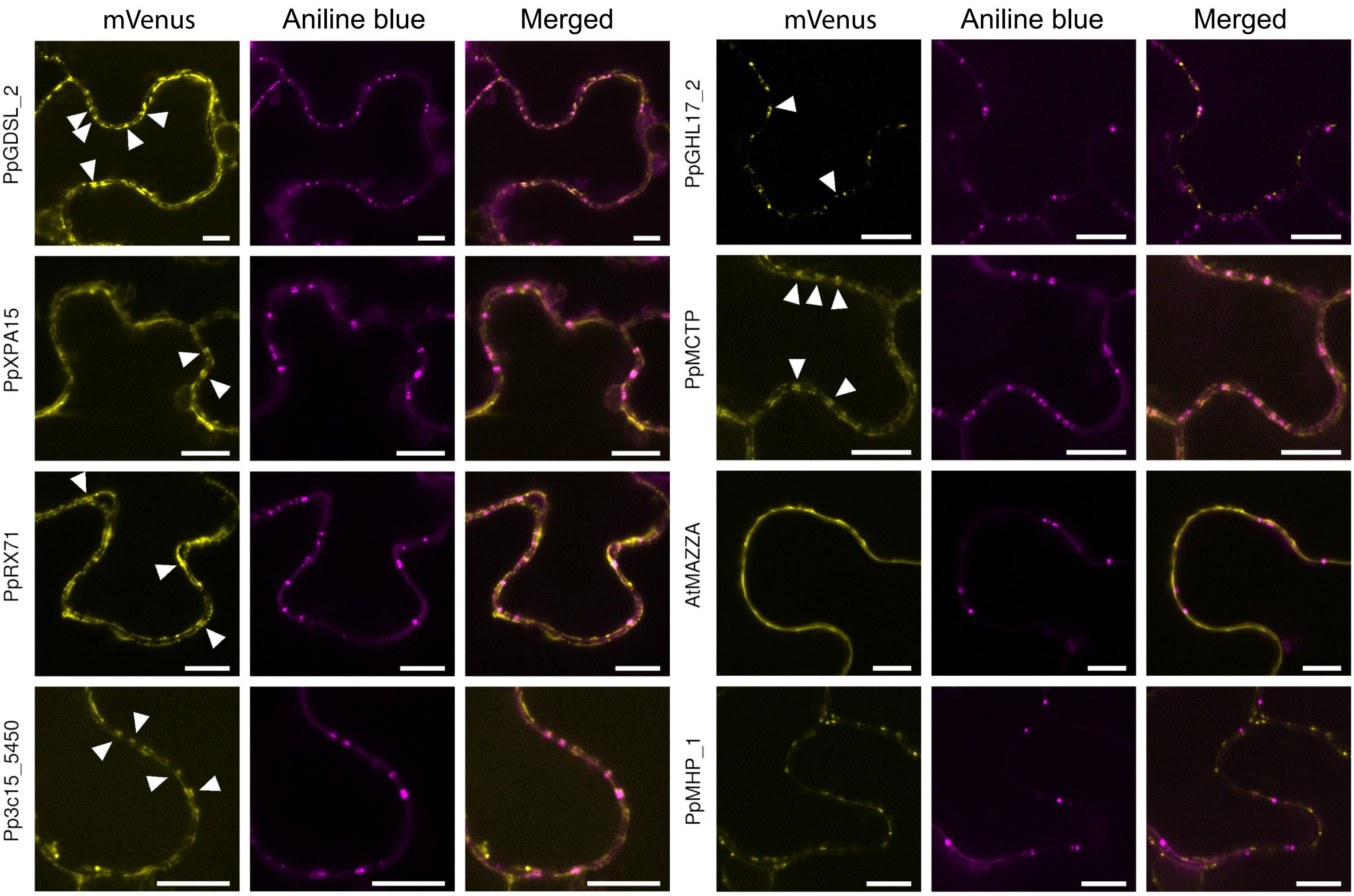


Supplementary Figure S5

### Supplementary Figure S6_GHL17s full

A

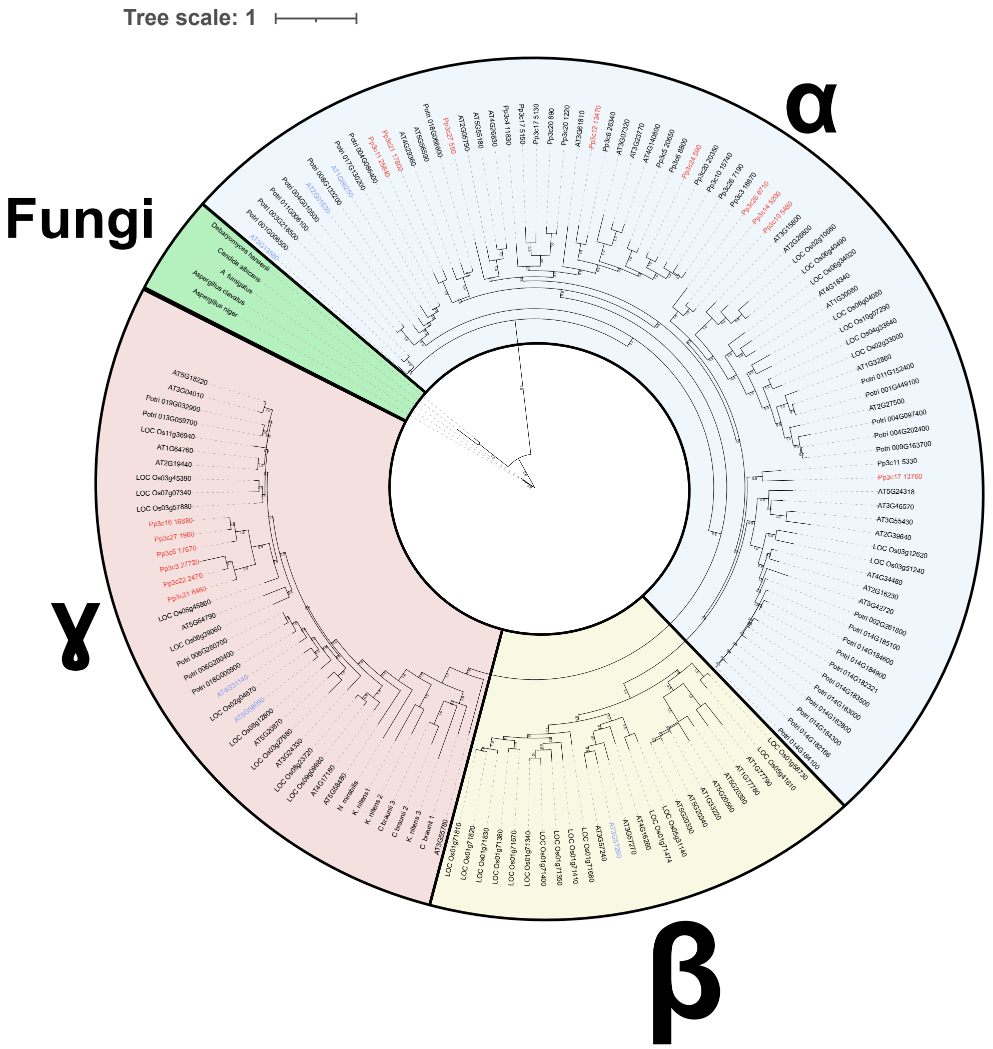


B

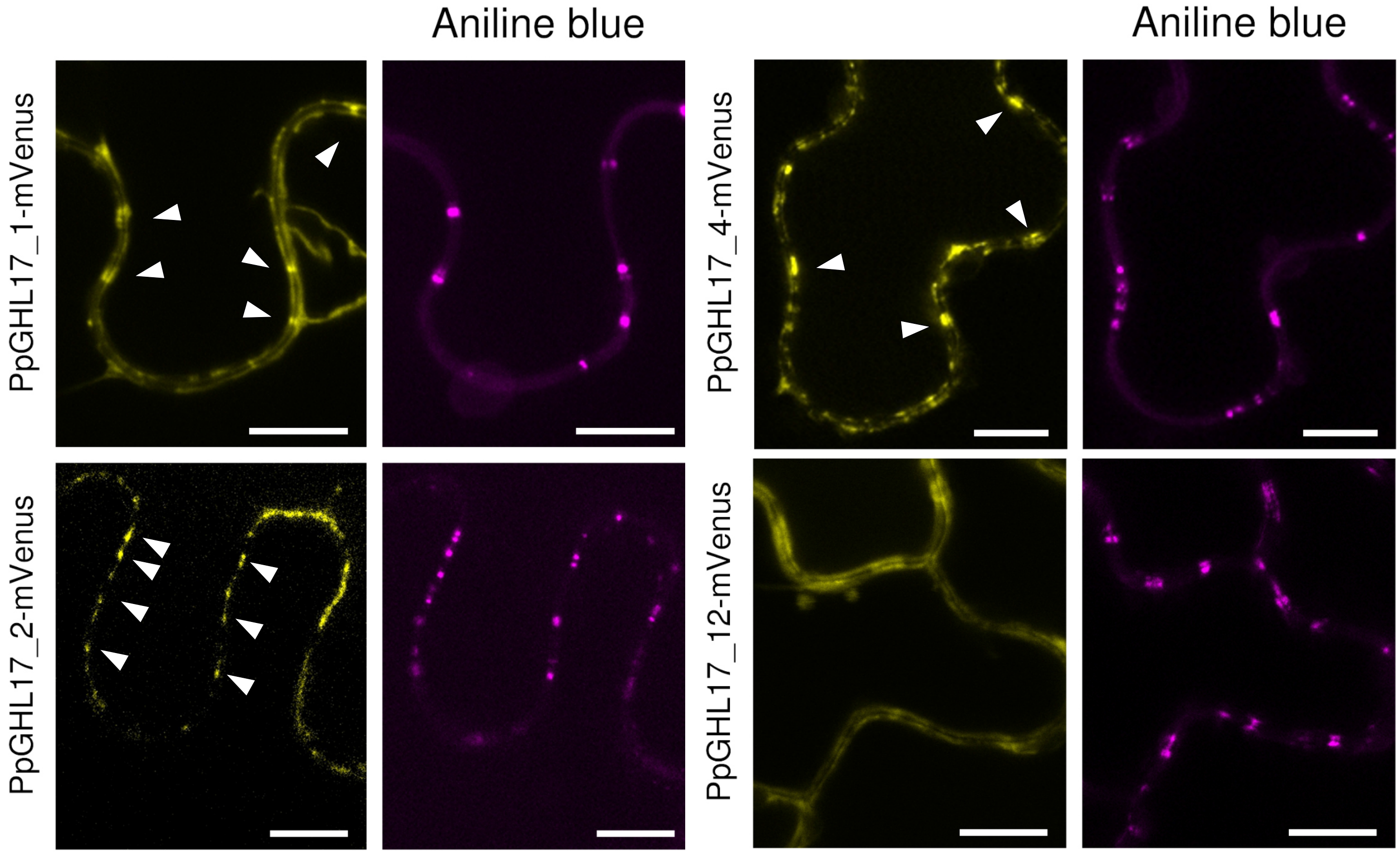


Supplementary Figure S6

### Supplementary Figure S7_XTHs

A

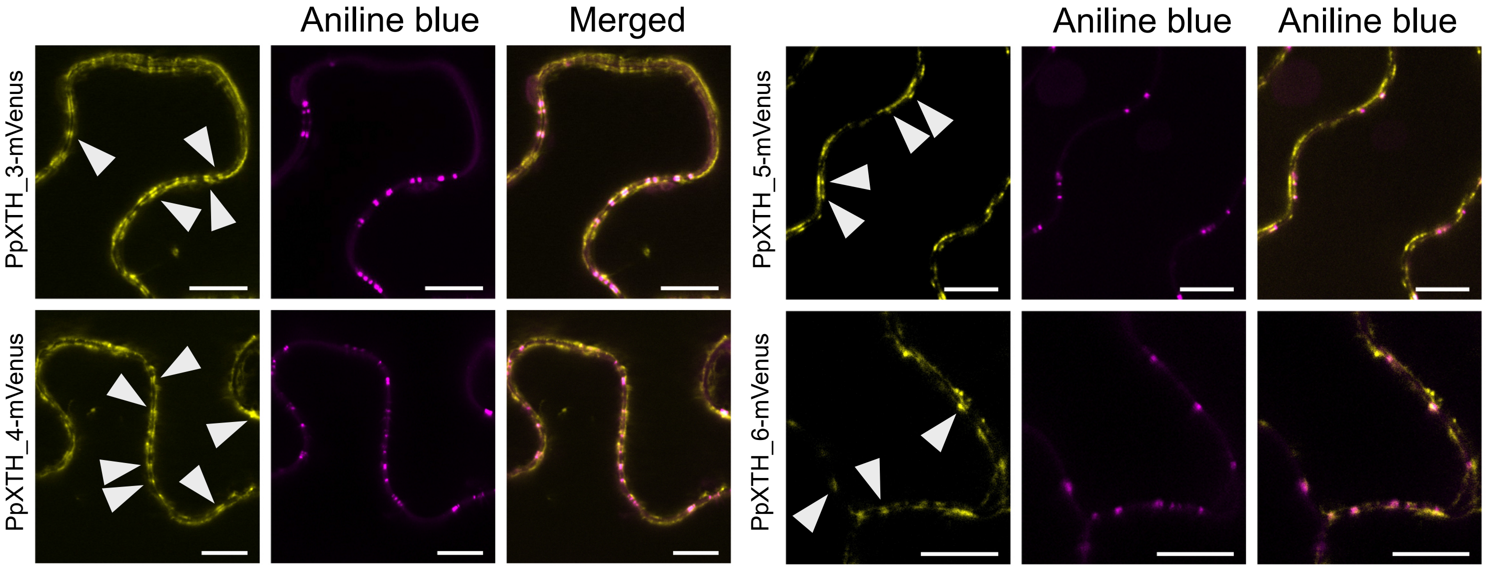


B

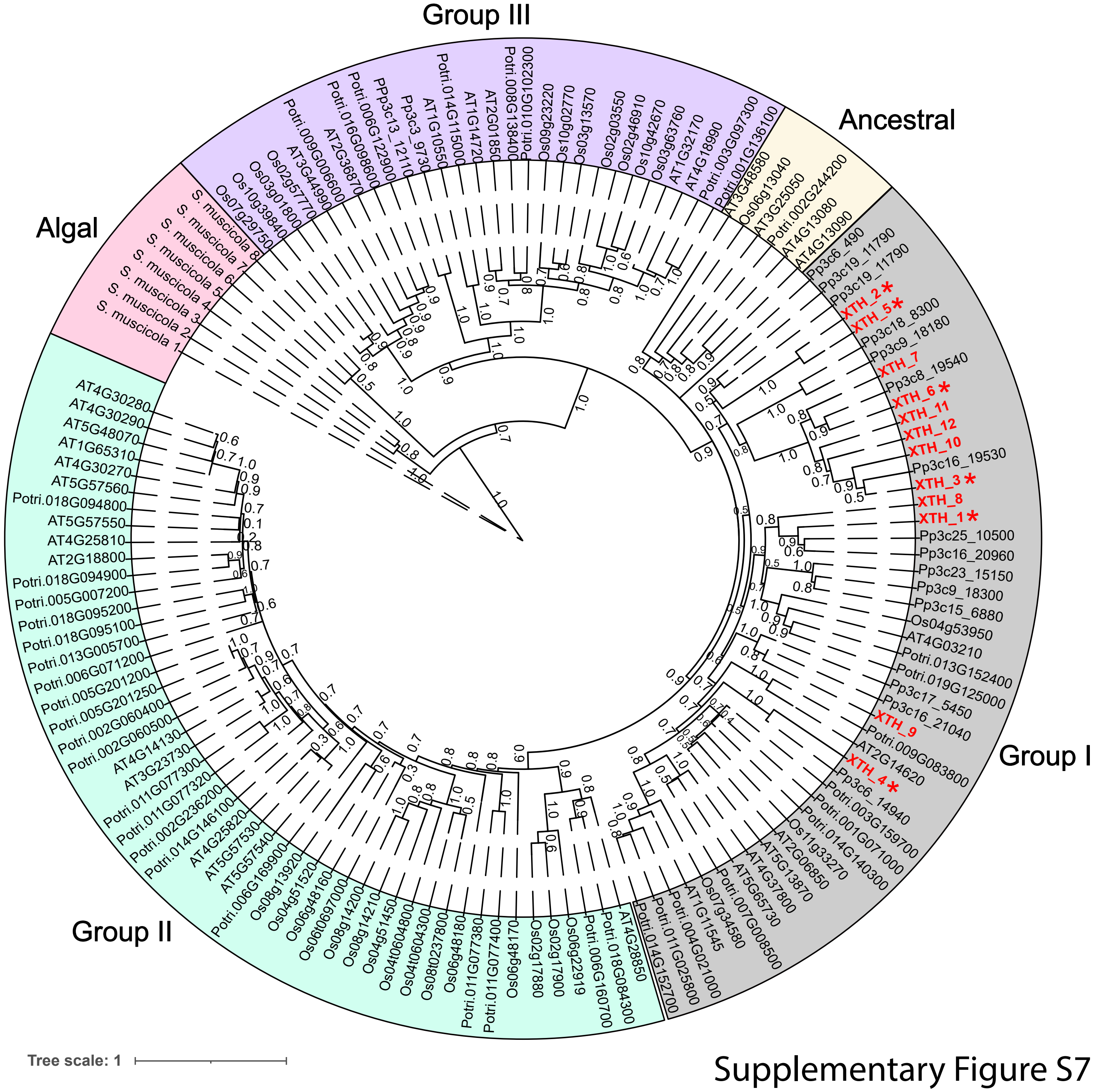


Supplementary Figure S7

### Supplementary Figure S8_EXO

**A**

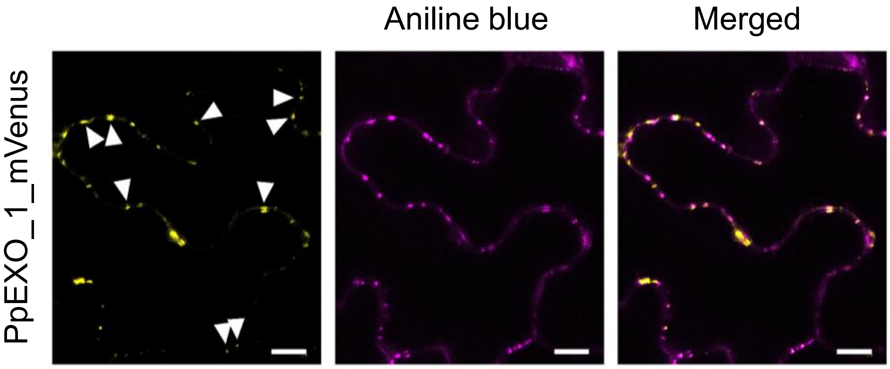


**B**

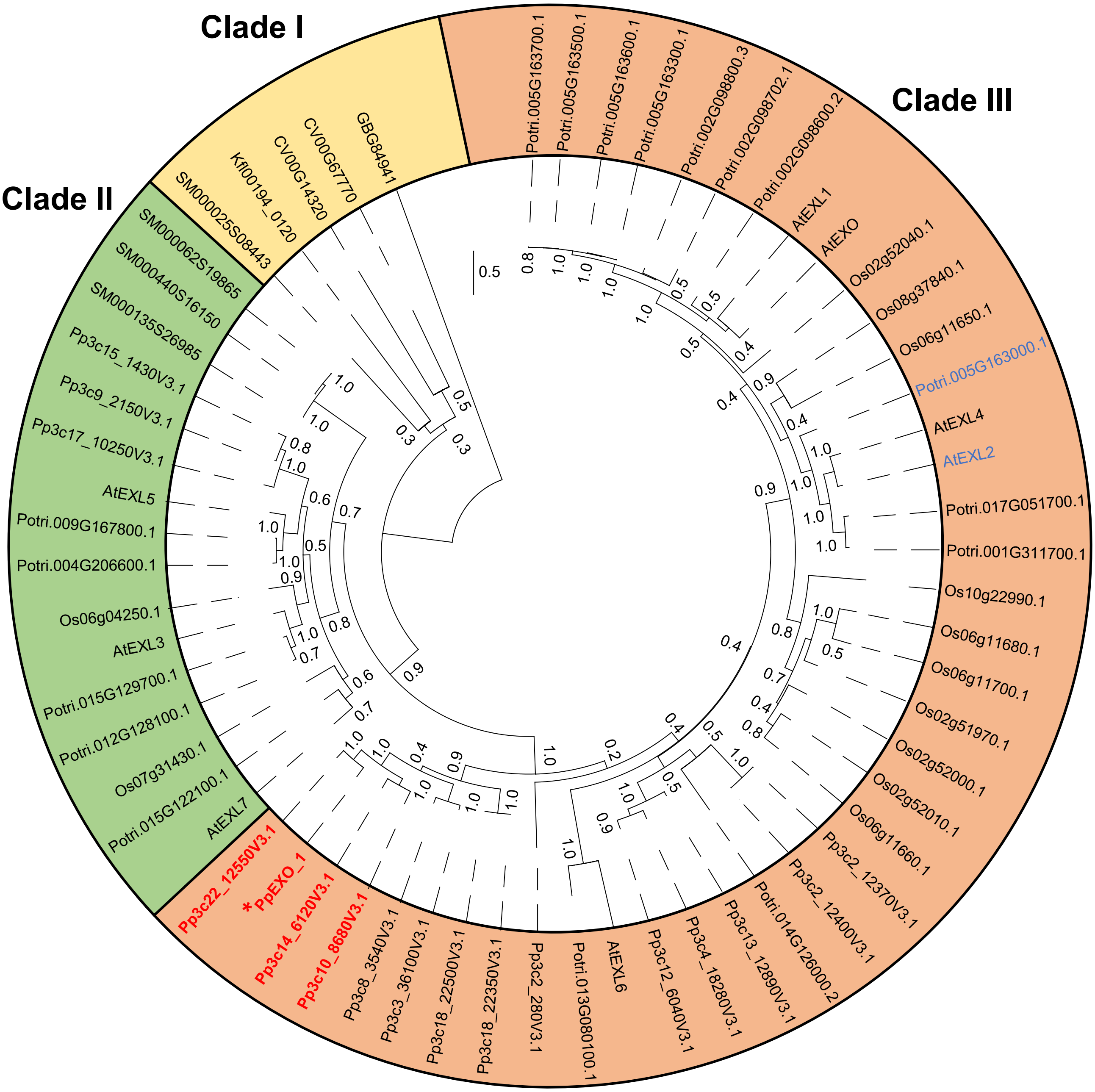


Supplementary Figure S8

### Supplementary Figure S9_venn diagrams

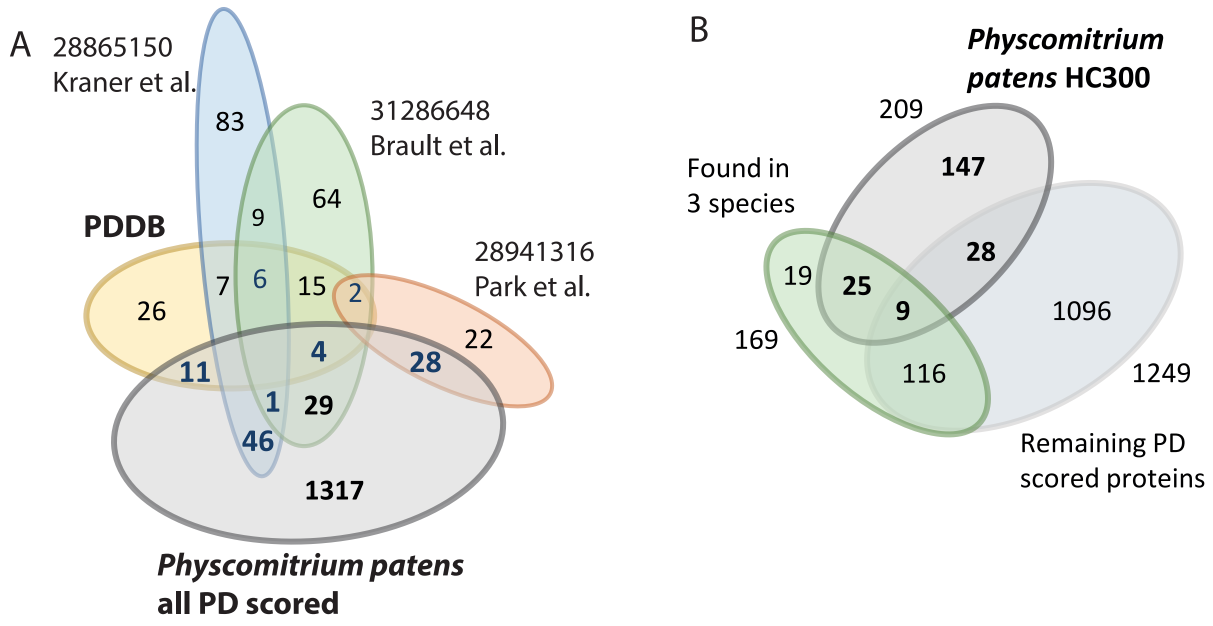


Supplementary Figure S9
